# Learning to Pursue Goals that Conflict with Prior Experience Relies on Fast Revaluation and Slow Value Updating

**DOI:** 10.64898/2026.10.16.759732

**Authors:** Ti-Fen Pan, Gaia Molinaro, Anne G.E. Collins

## Abstract

Humans can flexibly pursue novel goals, learning to reach them from goal-dependent teaching signals. However, the breadth of this flexibility and its underlying mechanisms remain unclear. For example, can humans flexibly learn to reach a typically avoided outcome (e.g., +0) while avoiding a typically sought one (+1)? We hypothesized that humans are sufficiently flexible to learn from counterintuitive goals, but at a cost, and this ability is supported by complementary temporal dynamics of rapid cognitive control and slower, iterative outcome value updating. In two experiments, we confirmed that participants learned when pursuing counterintuitive goals, but less effectively than standard goals. Furthermore, extended practice reduced performance gaps. To characterize the underlying mechanisms, we developed a novel computational model incorporating fast goal switching and slow outcome value updating. This model best captured group-level behavior; individuals differed in the efficiency of these processes, capturing differences in the cost initially incurred by counterintuitive goals. Crucially, a stronger initial cost did not permanently limit performance: slower value updating compensated for early inefficiencies in rapid adaptation. Furthermore, comparing early and late goal-switch learning revealed that greater goal alignment was associated with faster outcome processing, suggesting a transition from effortful control to efficient outcome evaluation. Thus, despite individual differences in initial flexibility, repeated attempts to reach counterintuitive goals decreased initial biases. More broadly, our findings show that goal-dependent outcome valuation leverages complementary fast and slow processes to enable flexible and efficient goal pursuit over time.

## Introduction

Humans exhibit an immense capacity to pursue novel and sometimes counterintuitive goals, such as rock climbing without ropes (free solo) or restricting calorie intake against ingrained survival instincts. Prior work has shown that humans are flexible in learning from abstract and novel goal outcomes (McDougle et al., 2022; Molinaro & Collins, 2023, 2026), with neuroimaging evidence highlighting how the prefrontal cortex (PFC) dynamically updates neural representations of outcome values given current goals (Frömer et al., 2019; Castegnetti et al., 2021).

Beyond abstract and novel goals, converging evidence indicates that humans can flexibly remap outcome values under counterintuitive goal conditions (e.g., minimizing gains; Shi et al., 2026) and learn from goal outcomes that run counter to prior experience (e.g., pursuing +0 over +1; Pan et al., 2025). Brain imaging studies have attributed this flexibility to support from multiple regions, including PFC, the reward network (McDougle et al., 2022; Hjort et al., 2026; Koszeghy & Passecker, 2026), and the hippocampus (Edelson & Hare, 2023; Chericoni et al., 2026). Despite this flexibility, pursuing counterintuitive goals often reduces performance, reflecting persistent influence from prior experience (Pan et al., 2025; Shi et al., 2026). Yet, the cause of this impairment remains unclear. In particular, we do not know whether this impairment reflects a fixed cost of processing counterintuitive goals by recruiting executive function, or whether this cost is temporary and one that practice can mitigate. This gap more generally reflects our poor understanding of what computational mechanisms underlie humans’ goal-dependent learning flexibility (Molinaro & Collins, 2023).

To address this gap, we used goal-dependent learning tasks to test whether practice can overcome learning inefficiency imposed by prior experience, and how distinct computational processes dynamically support goal-congruent value update. Drawing on hierarchical frameworks of goal-directed behavior (O’Reilly, 2020; Koszeghy & Passecker, 2026; Xu et al., 2026), we hypothesized that goal-dependent valuation and learning recruits two complementary mechanisms: a fast and effortful process that governs rapid goal-dependent outcome evaluation, and a slow process that trains a goal-congruent outcome-reward association (i.e., an association between the current goal outcomes and the positive reward associated with them; Molinaro and Collins, 2026). Specifically, in this framework, the fast process depends on the active goal representation maintained in executive control, which modulates the slow, gradual updating of goal-congruent mappings while remaining constrained by prior experience.

To test the dynamics of the goal-dependent valuation framework, we first designed a self-determined goal task in which participants determined their own goals in a stimulus-action learning task, establishing baseline behavioral flexibility and prior bias across standard versus counterintuitive goal outcomes (e.g., pursuing +1 vs. +0; green vs. red outcomes). To dissect the dynamics of goal-dependent learning, we then implemented an extended stimulus-action learning task involving explicit, instructed goal switches between opposing goal outcomes.

Our behavioral results showed that participants initially pursued standard outcomes (+1 over +0), but extended practice enabled flexible adaptation to counterintuitive goals (pursuing +0) and eliminated the performance gap between goals. Computational modeling through a goal-dependent reinforcement learning (RL) framework revealed different degrees of efficiency in fast and slow valuation mechanisms, leading to individual variation in susceptibility to prior experience. Over the time course of learning, practice attenuated these prior constraints through complementary fast and slow valuations, driving a transition from effortful, controlled processing to flexible, automated goal execution.

## Results

### Prior experience shapes the attribution of outcome-reward association

#### Experiment 1

We first designed a goal-dependent stimulus-action learning task to investigate whether humans select different goals during learning based on prior experience, and how these prior preferences impact learning when explicit goals align with or contradict them. Participants completed two sequential blocks: an initial self-determined block followed by an instructed block. Each block consisted of 26 trials comprising 13 iterations of two distinct visual stimuli, with an entirely novel stimulus set introduced in each block.

Across both blocks, the core task followed a deterministic stimulus–action learning paradigm (Fig. 1A). On each trial, participants observed a stimulus, selected one of three key presses, and received deterministic outcome feedback (Fig. 1 A). In the instructed block (the second block), the goal was instructed and remained fixed throughout the block. On each trial, participants were first informed of which outcome (+1 or +0) was their goal, and were instructed to select actions leading to the goal outcome. By contrast, in the self-determined block (the first block), participants were not told which goal outcome to pursue, only what the possible outcome values were (+1 or +0); the stimulus-action outcome remained deterministic based on the experimenter-defined goal (unknown to the participants). For example, in a Goal 0 self-determined goal, one correct action (e.g., J for the mushroom in Supplementary Fig 6) systematically led to the goal outcome (e.g., +0), while the other two actions led to the non-goal outcome (e.g., +1). Participants were told that instructions were lost and to try to play the game (see Methods for more details). Administering the self-determined block first allowed us to capture participants’ initial biases toward desirable goal outcomes while precluding carryover effects from explicit instruction. After completing the self-determined block, participants rated their preference for each outcome from 0 to 100 and indicated the strategy they used during the task (e.g., collecting as many +1 as possible). They then proceeded to the second block, the instructed block, with the same goal but a new set of stimuli.

**Figure 1.**
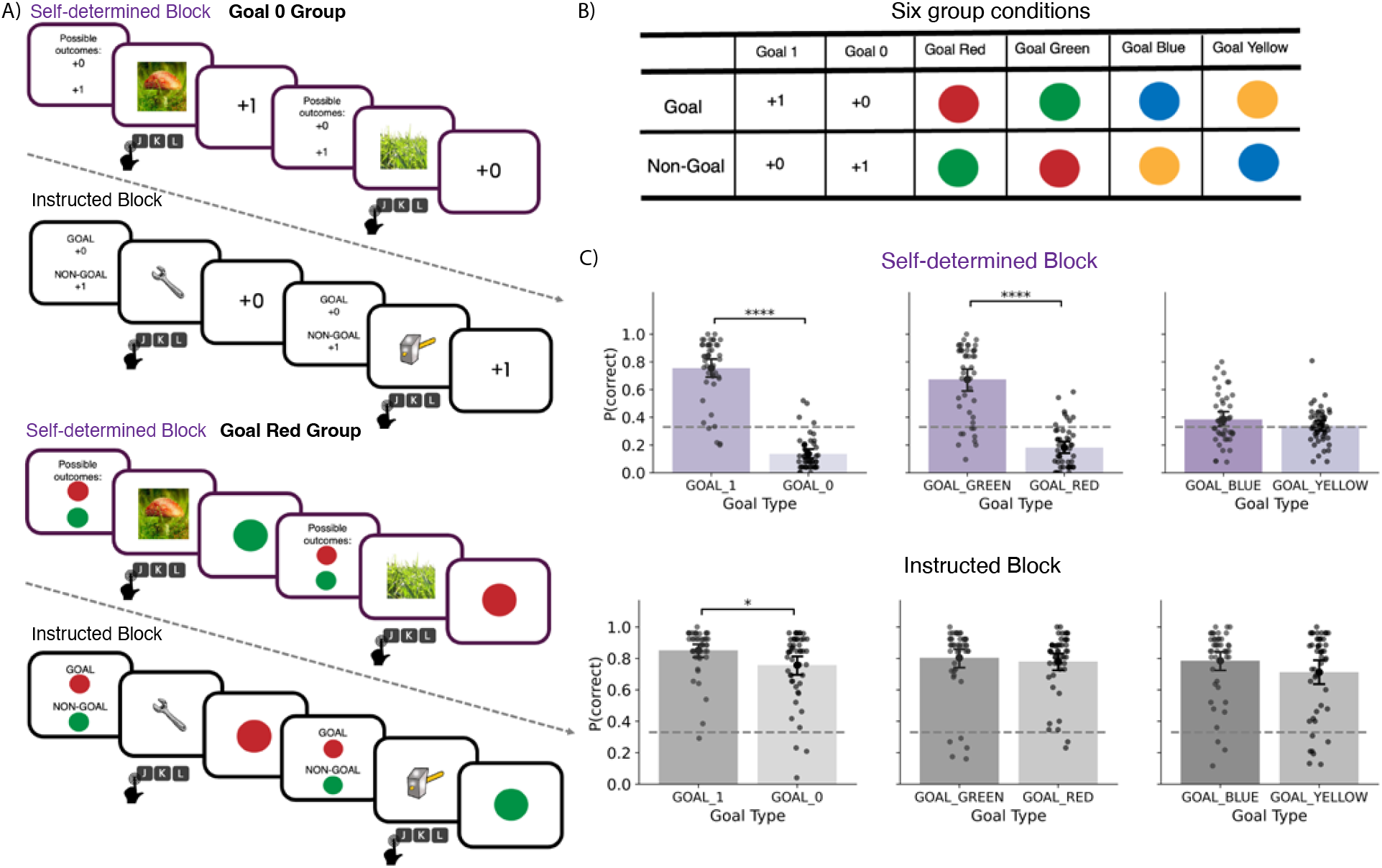
Experiment 1 paradigm and behavior results: **A)** The goal-dependent learning experiment comprised two short blocks: a self-determined and an instructed block. Top: Goal 0 group (Goal:+0, Non-Goal:+1). Bottom: Goal Red group (Goal: Red, Non-Goal: Green). **B)** Goal and non-goal outcomes of six different groups. **C)** Average choice accuracy across goal groups in the self-determined block (top) and instructed block (bottom). Each dot is a single participant’s data. Error bars indicate 95% CI. Sample sizes for each group: Goal 1 (n = 46*), Goal 0 (n* = 49*), Goal Red (n* = 48*), Goal Green (n* = 46*), and Goal Blue (n* = 46). ^∗^*p <* 0.05; ^∗∗∗∗^*p <* 0.0001*; Unpaired two-sided t-test with Benjamini–Hochberg correction for multiple comparisons*.

To investigate different outcomes’ impact, we designed six different goal/non-goal pairs: +1/+0 (Goal 1), +0/+1 (Goal 0), green/red (Goal Green), red/green (Goal Red), blue/yellow (Goal Blue), yellow/blue (Goal Yellow). Each participant was randomly assigned to one of the groups. We expected that the Goal Blue and Goal Yellow groups would show no strong preference for outcomes and thus serve as control groups (Fig. 1 B); by contrast, given the majority Western cultural associations of our participant cohort, we anticipated that outcomes +1 and green would be strongly preferred in the absence of instructions.

#### Flexible learning with different goals

In the self-determined block, participants in the Goal 0 and Goal Red groups exhibited lower accuracy compared to the Goal 1 and Goal Green groups, respectively (Fig. 1 C; Goal 0 vs Goal 1: *t*(67.88) = −16.46, *p* < .001, Bayes factor (BF_10_) *>* 100; Goal Red vs Goal Green: *t*(70.42) = −10.90, *p* < .001, BF_10_ *>* 100). Specifically, Goal 0 and Goal Red groups systematically selected actions leading to non-goal outcomes, driving performance well below chance (one-sample *t*-tests against 0.33; Goal 0: *M* = 0.14, *t*(48) = −11.22, *p* < .001, BF_10_ = 1.33 × 10^12^, 95% CI [0.10, 0.17]; Goal Red: *M* = 0.18, *t*(47) = −6.78, *p* < .001, BF_10_ = 7.27 × 10^5^, 95% CI [0.14, 0.23]). This systematic bias suggests that participants viewed 1 and Green as the “correct” outcome to pursue based on prior experience. Self-reported strategies corroborated this finding: participants in Goal 0 and Goal Red groups frequently reported collecting non-goal outcomes (+1 and Green; Supplementary Fig. 7 A). Furthermore, these self-determined rules strongly correlated with participants’ subjective preferences for different outcomes, as indicated by their post-block ratings (Supplementary Fig. 7 B-D).

As expected, participants in the Goal Blue and Goal Yellow groups, lacking strong preferences given past experience, exhibited more dispersed behavior, achieving around chance-level accuracy (33%) in the self-determined block. Moreover, we found no difference in accuracy between these goals (Goal Yellow vs Goal Blue: *t*(81.91) = −1.42, *p* = .159, BF_10_ = 0.53, with median posterior *δ* = 0.26, 95% CI [−0.12, 0.65]). Notably, few participants reported collecting rare outcomes, which corresponded to the correct rule in our task (Supplementary Fig. 7 A). Discovering this strategy requires extensive trial-and-error learning and is considerably more cognitively demanding than relying on standard rewarding outcomes guided by intuitive bias.

Having confirmed that participants came into the experiment with strong priors for +1/Green as desirable goals and +0/Red as outcomes to avoid, we next examined how flexibly participants could override those priors. In the full instruction block, participants achieved high average accuracies ranging from 71.2% to 85.1% across all goal types (Fig. 1 C). Crucially, performance in every condition was significantly above the 33% chance level (one-sample *t*-tests: all *t*s ≥ 9.88, all degrees of freedom between 45 and 48, all *p*_adj_ *< .*001,], all BF_10_ ≥ 1.70 × 10^10^, 95% CIs within [0.63, 0.89]). These results showed that humans can flexibly learn to pursue arbitrary and even counterintuitive goal outcomes when given clear instructions.

#### Learning inefficiency in Goal 0

Despite the behavioral flexibility observed in the instructed block, participants exhibited attenuated learning performance in Goal 0 compared to Goal 1 (*t*(84.31) = −2.47, *p* = .016, BF_10_ = 2.879, with median posterior *δ* = 0.451, 95% CI [0.06, 0.854]). By contrast, no significant performance differences emerged between color-pair goals (Goal Red vs. Goal Green, or Goal Yellow vs. Goal Blue). This is particularly notable in the case of the Red group, which exhibited a pronounced prior against Red outcomes observed in the self-determined block.

This asymmetry between Goal 0 and Goal Red suggests that intuitive priors for numerical magnitude are stronger and more uniform across individuals than those for color outcomes. While nearly everyone prefers a +1 outcome to a +0, color preference is more subjective and culturally dependent, leading to more individual differences (e.g., some individuals may prefer red over green; Supplementary Fig. 7 B). This pattern was further supported by a linear mixed-effects model (LMM) predicting accuracy from block (self-determined vs. instructed), pair type (color vs. numerical), goal counterintuitiveness (Goal 0 and Goal Red coded as counterintuitive), and their interactions. While the three-way interaction was not significant, the model revealed a significant interaction between goal counterintuitiveness and pair type (*β* = 0.094, *SE* = 0.044, *t*(373.81) = 2.158, *p* = .032; Supplementary Table 2) alongside an interaction between counterintuitiveness and block (*β* = 0.493, *SE* = 0.03, *t*(373.81) = 16.446, *p* < .001). Accuracy was higher for numerical than color pairs under standard goals (Goal 1 vs. Goal Green), whereas the pattern was reversed under counterintuitive goals (Goal 0 vs. Goal Red), indicating a stronger prior bias for numerical over color outcomes across blocks.

In addition, we conducted a control experiment to assess potential practice effects between blocks, particularly for conventional goals such as Goal +1 and Goal Green. We found that practice effect alone cannot account for the performance differences across blocks, suggesting that our instructional manipulation drove the observed learning differences. For further details, see Supplementary Fig. 8.

Together, Experiment 1 showed that without explicit rules, prior experience steered most participants to pursue conventional rewarding outcomes (e.g., +1, green). However, under explicit instructions, they were able to flexibly learn from counterintuitive goals (e.g., +0, red). Nonetheless, this flexibility could be costly, as participants learned less effectively from Goal 0 relative to Goal 1. This performance deficit likely reflects the heightened demand placed on limited executive control resources to dynamically update outcome values (Molinaro & Collins, 2026; Shi et al., 2026). While these results confirmed a fast dynamic for goal value setting, a key question remains: is there also a slower dynamic for outcome-value update, and can extended practice overwrite the influence of prior experiences?

### Long-term dynamics of goal-dependent valuation and learning

#### Experiment 2

Experiment 1 showed that humans have strong pre-existing outcome-reward associations and can override them in a goal-dependent way at a cost, showing evidence for a rapid outcome valuation process. In Experiment 2, we examined whether a slower outcome evaluation process also exists, such that practice decreases the learning inefficiency that the rapid process cannot overcome. To do so, we designed a goal-dependent learning task with both slow and fast dynamics of changes in goals. As in Experiment 1, participants learned deterministic stimulus-action mappings. Unlike Experiment 1, participants were always explicitly instructed which outcome to pursue. Additionally, to assess outcome processing efficiency, we introduced a Go/No-go “collect” mechanism: participants were instructed to use a separate key press to *collect* target outcomes and to ignore non-goal outcomes (Fig. 2A). For counter-intuitive goals, we used Goal 0, informed by the results of Experiment 1, where the Goal 1 and Goal 0 conditions exhibited the largest performance contrast.

**Figure 2.**
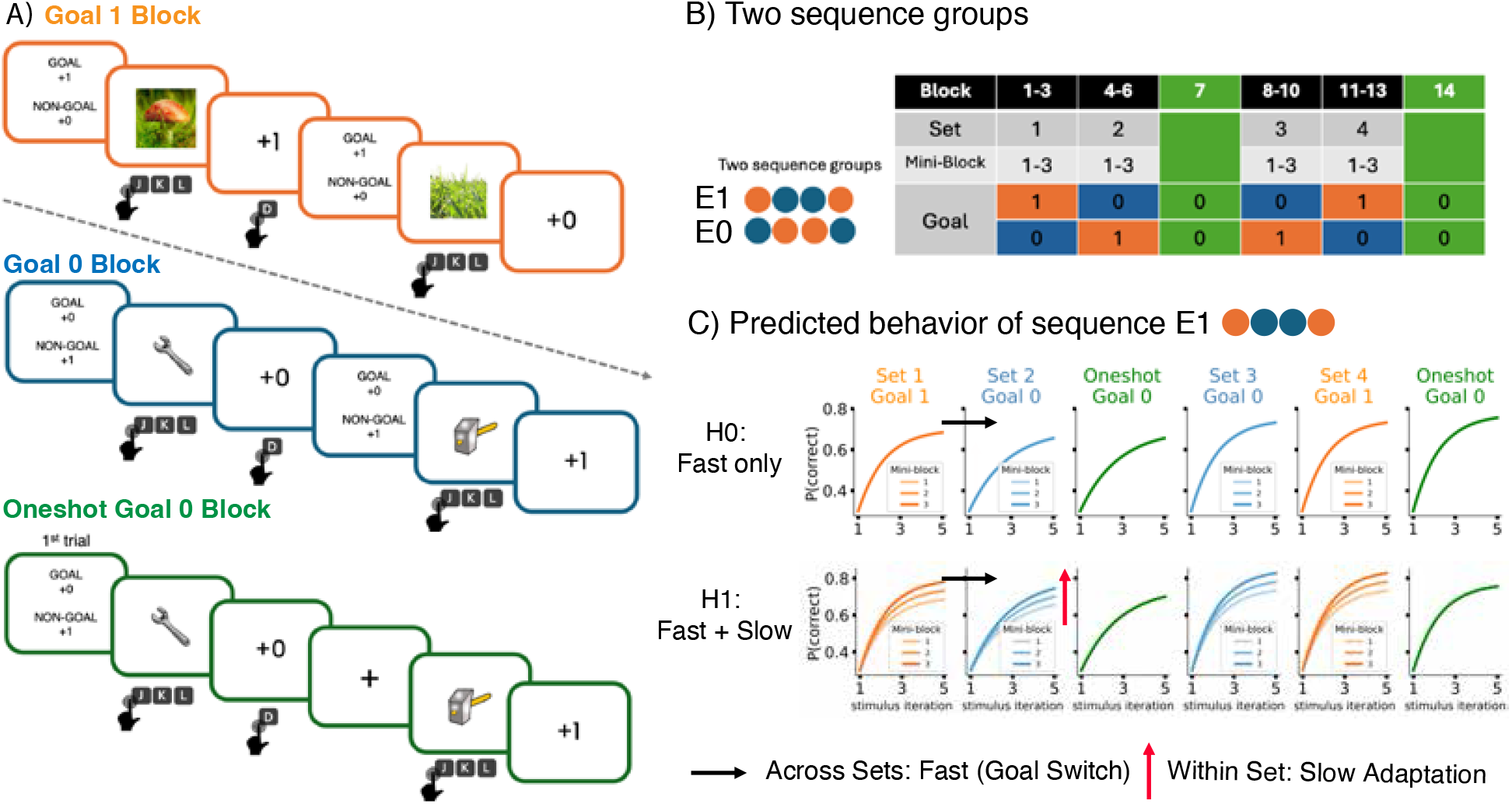
Experiment 2 paradigm and predicted dynamics. **A)** Goal-dependent learning task with explicitly instructed goals. **B)** Task structure for the two goal sequence groups: Group E1 (goal sequence: 111, 000, 0, 000, 111, 0) and Group E0 (goal sequence: 000, 111, 0, 111, 000, 0). **C)** Predicted behavioral dynamics for sequence group E1. Hypothesis 0: fast only, a fast goal switch across block sets (e.g., Set 1 - Block 3 to Set 2 - Block 4). Hypothesis 1: fast+slow, fast alongside slow reward adaptation within a block set across mini-blocks (e.g., Blocks 4 to 6).

The experiment comprised 14 blocks across three distinct conditions: Goal 1, Goal 0, and one-shot Goal 0. In Goal 1 and Goal 0 blocks, the designated target outcome was displayed at the onset of each trial and remained constant throughout the block. By contrast, during one-shot Goal 0 blocks, the goal (+0) and non-goal (+1) reminders were presented only on the initial trial, requiring participants to maintain the active goal representation across the entire block. Within each block, participants learned new mappings between 6 novel stimuli and 3 actions, with 5 iterations per stimulus (30 trials per block). This brief block structure kept participants in an active learning regime where outcomes remained informative, enabling us to isolate how shifting goals affect new learning.

To dissociate fast and slow dynamics of goal-dependent valuation, we structured the block order using alternating runs of repeated and switched goals across two between-subject groups (Fig. 2 B). A “set” was defined as three consecutive blocks sharing the same goal, with individual constituent blocks serving as mini-blocks. Participants were randomly assigned to one of two sequence structures consisting of these block types. The E1 group (Early 1 Late 0) began with three consecutive Goal 1 blocks (Set 1; Blocks 1–3) before switching to three consecutive Goal 0 blocks (Set 2; Blocks 4–6). Following a one-shot Goal 0 block (Block 7), the E1 group completed another three Goal 0 blocks (Set 3; Blocks 8–10), three Goal 1 blocks (Set 4; Blocks 11–13), and a final one-shot Goal 0 block (Block 14). Following a parallel structure, the E0 group experienced an inverted set sequence (Goal 0, Goal 1, Goal 1, and Goal 0), with one-shot Goal 0 blocks likewise at Blocks 7 and 14.

These sequence manipulations were designed to isolate fast (at goal-switch blocks) from slow dynamics (across repeated goal blocks, and the entire task; Fig. 2C). Based on the rapid adaptation observed in Experiment 1, we predicted that participants would quickly update their goal representations across sets to learn from feedback (fast dynamics), with a cost to counterintuitive goals. Concurrently, if present, slow dynamics would manifest as a gradual updating of the outcomes’ intrinsic reward properties within sets (across mini-blocks), thereby improving overall learning efficiency (Fig. 2C, H0 vs. H1). Furthermore, we expected one-shot blocks to provide more sensitivity to slow dynamics, with only one opportunity to apply task instructions rapidly. Consequently, performance in the late one-shot Goal 0 block would surpass that of the early one-shot block, driven by progressively stabilized goal-congruent outcome–reward associations. To test these hypotheses, we analyzed stimulus–action learning at both the trial and block levels. We focused specifically on goal-switch sets (Sets 2 and 4) to evaluate the learning costs of goal-switching and the influence of prior experience. Additionally, we measured reward collection (RC) precision (d’) and response times (RT) to determine how practice and prior experience affect the accuracy and speed of goal-dependent outcome processing.

#### Initial learning inefficiency in Goal 0

We first analyzed learning in the initial block (Block 1) to replicate Experiment 1’s results, showing successful but impaired Goal 0 learning. As expected, both groups performed significantly above chance (33%): participants in the E1 group achieved a mean accuracy of 60.2% (95% CI [56.0%, 64.0%], *t*(35) = 12.97, *p* < .001, BF_10_ = 1.07 × 10^12^; two-sided one-sample *t*-test), and participants in the E0 group achieved 54.6% (95% CI [50.0%, 59.0%], *t*(39) = 9.16, *p* < .001, BF_10_ = 3.31 × 10^8^). A generalized linear mixed-effects model (GLMM) predicting trial-level accuracy from sequence group and stimulus iteration revealed a positive learning slope across iterations (*b* = 1.06, *SE* = 0.06, *z* = 17.39, *p* < .001, OR = 2.88, 95% CI [2.56, 3.25]; Supplementary Table 3). Critically, the group under Goal 0 exhibited significantly lower overall accuracy than those in the Goal 1 condition (*b* = −1.20, *SE* = 0.10, *z* = −11.51, *p* < .001, OR = 0.30, 95% CI [0.25, 0.37]). Block 1 results thus replicate Experiment 1 findings: learning efficiency was lower for Goal 0 than for Goal 1 in the initial learning block.

#### Early goal-switch block reveals fast valuation

To further validate the flexibility of goal-dependent learning, we examined the first goal-switch block (Block 4). We predicted that both groups would adapt to the new goal, but that switching to Goal 0 would incur a higher cognitive cost. Analysis of Block 4 confirmed that participants demonstrated fast and flexible goal-switching, achieving above-chance accuracy whether transitioning to Goal 0 or Goal 1. Both sequence groups performed significantly above chance (33.3%) in Block 4, confirming rapid goal switching: the E0 group (switching to Goal 1) achieved a mean accuracy of 53.1% (95% CI [46%, 60%], *t*(39) = 5.52, *p* < .001, BF_10_ = 7440.63; two-sided one-sample *t*-test), and the E1 group (switching to Goal 0) achieved 51.1% (95% CI [44%, 58%], *t*(35) = 5.10, *p* < .001, BF_10_ = 1765.20). Next, to confirm that switch cost exists in both groups, we compared accuracy between the pre-switch block (Block 3) and the post-switch block (Block 4) within each group.

We found significant drops in accuracy for both the E1 group (switching to Goal 0: *t*(35) = 4.41, *p* < .001, 95% CI [0.08, 0.21], BF_10_ = 263.73; two-sided paired *t*-test) and the E0 group (switching to Goal 1: *t*(39) = 3.38, *p* = .005, 95% CI [0.05, 0.21], BF_10_ = 19.50). Contrary to our prediction, a two-sided unpaired *t*-test showed no significant difference in accuracy drops between the groups. Thus, reconfiguring outcome valuation imposed a substantial cost, regardless of whether participants switched to Goal 0 or 1. Lastly, to further examine the difference in switch costs between groups, we fitted an analogous GLMM to Block 4 trial-level accuracy.

Consistent with the lack of switch cost difference, there was no effect of group, but only a main effect of stimulus iterations (*b* = 0.92, *SE* = 0.09, *z* = 10.72, *p* < .001, OR = 2.50, 95% CI [2.12, 2.96]; Supplementary Table 4). However, a significant two-way interaction suggests that participants switching to Goal 1 (E0) exhibited a steeper learning slope than those switching to Goal 0 (E1) (*b* = 0.37, *SE* = 0.17, *z* = 2.14, *p* = .032, OR = 1.44, 95% CI [1.03, 2.01]). This indicates that learning under Goal 0 in Block 4 did exhibit a learning deficit, although its effect was too weak to significantly decrease overall accuracy. Together, the above-chance performance points to a fast valuation mechanism capable of rapidly adapting to changing goals. However, the substantial switch costs observed across both goals led us to ask whether a complementary, slower valuation mechanism also exists—one that makes the typically rewarding pursuit of Goal 1 less efficient after sustained exposure to Goal 0.

#### Early block sets reveal slow valuation

In our sequence manipulation, participants learned the same goals across consecutive blocks, which we refer to as “mini-blocks”. We hypothesized that learning efficiency would incrementally improve across these mini-blocks. To test whether a slow process contributes to goal outcome valuation, we fitted an LMM model predicting accuracy from sequence group and mini-block in Set 1 and Set 2. There was no main effect of the sequence group nor a two-way interaction, however, the main effects of mini-blocks in both sets indicated a slow improvement across mini-blocks (Set 1: *b* = 0.042, *SE* = 0.010, *t* = 4.31, *p* < .001; Set 2: *b* = 0.070, *SE* = 0.012, *t* = 6.04, *p* < .001; Supplementary Table 5& 6). Specifically, in the goal-switch set (Set 2), this improvement across mini-blocks suggests the presence of a slower valuation mechanism that complements the fast valuation, regardless of whether participants switched to Goal 0 or Goal 1. Although accuracy was higher under Goal 1 (61.5%) than under Goal 0 (58.2%) in Set 2, the lack of a significant difference indicates that the initial learning impairment associated with Goal 0 decreases as slow valuation unfolds over consecutive blocks. Note that this cannot be interpreted as an effect of time on task, as performance drops in Block 4 from Block 3 when goals switch.

#### Extended practice reduces goal-switch cost

Our early-learning findings suggest that goal-dependent learning reflects complementary fast and slow valuation mechanisms: participants rapidly adapted to changing goals, while learning efficiency improved across consecutive mini-blocks. However, the high switch cost observed in early learning prompted us to test whether in later goal-switch blocks (Blocks 8 and 11), continued practice enhances learning efficiency when facing changing goals. In Block 8, following a one-shot Goal 0 block (Block 7), the E0 group switched goals (learning under Goal 1), whereas the E1 group continued learning under Goal 0. Indeed, a trial-level GLMM predicting accuracy from sequence group, stimulus iteration, and their interaction revealed main effects of iteration (*b* = 1.70, *SE* = 0.09, *z* = 17.99, *p* < .001, OR = 5.47, 95% CI [4.55, 6.58]; Supplementary Table 7), sequence group (*b* = −0.64, *SE* = 0.18, *z* = −3.52, *p* < .001, OR = 0.53, 95% CI [0.37, 0.75]), and two-way interactions (*b* = −0.58, *SE*= 0.19, *z* = −3.15, *p* = .002, OR = 0.56, 95% CI [0.39, 0.80]), indicating the consequences of a goal switch on learning performance.

By Block 11, however, we found no evidence of a switch cost in either group. Within-group comparison of accuracy between the pre-switch block (Block 10) and the post-switch block (Block 11) showed no significant decrease. Furthermore, we observed a significantly greater drop in accuracy during the early switch (Blocks 3 to 4) compared to the late switch (Blocks 10 to 11) in both the E1 group (*t*(35) = −3.07, *p* = .008, 95% CI [−0.20, −0.04], BF_10_ = 9.11) and the E0 group (*t*(39) = −2.98, *p* = .008, 95% CI [−0.23, −0.04], BF_10_ = 7.51; two-sided paired *t*-tests). These results suggest that a stable goal-congruent outcome-reward association was trained to complement the fast goal switch. Despite the reduced switch cost, we found a performance difference between sequence groups at Block 11 (main effect of sequence group in GLMM: *b* = 0.46, *SE* = 0.18, *z* = 2.54, *p* = .011, OR = 1.58, 95% CI [1.11, 2.25]; Supplementary Table 8). We therefore examined the final block set (Set 4; Blocks 11–13) to determine whether slow valuation persists in reducing this between-group difference.

An LMM predicting accuracy from sequence group, mini-block, and their interaction revealed a significant effect of mini-block (*b* = 0.018, *SE* = 0.008, *t* = 2.12, *p* = .034, 95% CI [0.001, 0.035]; Supplementary Table 9), indicating that accuracy improved across mini-blocks in the late learning stage. Importantly, the main effect of sequence group was not significant, suggesting no overall difference in accuracy between groups. The significant two-way interaction indicated that the E1 group, learning under Goal 1, had a steeper increase in accuracy across mini-blocks than the E0 group, learning under Goal 0 (*b* = −0.051, *SE* = 0.017, *t* = −3.03, *p* = .002, 95% CI [0.018, 0.084]). This finding confirmed that slow valuation persists in the late learning stage, reducing the initial learning difference across groups.

Finally, in addition to evaluating adaptation across learning blocks, we assessed whether practice mitigated switch costs during one-shot Goal 0 blocks (Blocks 7 and 14), in which the goal reminder appeared only once at the beginning of a block and thus placed higher demands on internal goal maintenance. Within each sequence group, we tested the interaction between trial-level learning slopes and goal switch. For the E0 group, which experienced an early goal switch from Goal 1 to Goal 0, the interaction was significant (*β* = 0.38, *SE* = 0.16, *z* = 2.31, *p* = .021; Supplementary Table 10). This reflects a marked goal-switch cost during early learning that was absent at the late stage (Fig. 3A). By contrast, the E1 group, which underwent a late goal-switch, exhibited no such interaction (*β* = 0.02, *SE* = 0.18, *z* = 0.11, *p* = .911; Supplementary Table 11), showing stable learning slopes across stages. This confirms that practice accrued by the E1 group effectively counteracted the goal-switch cost at the later stage.

**Figure 3.**
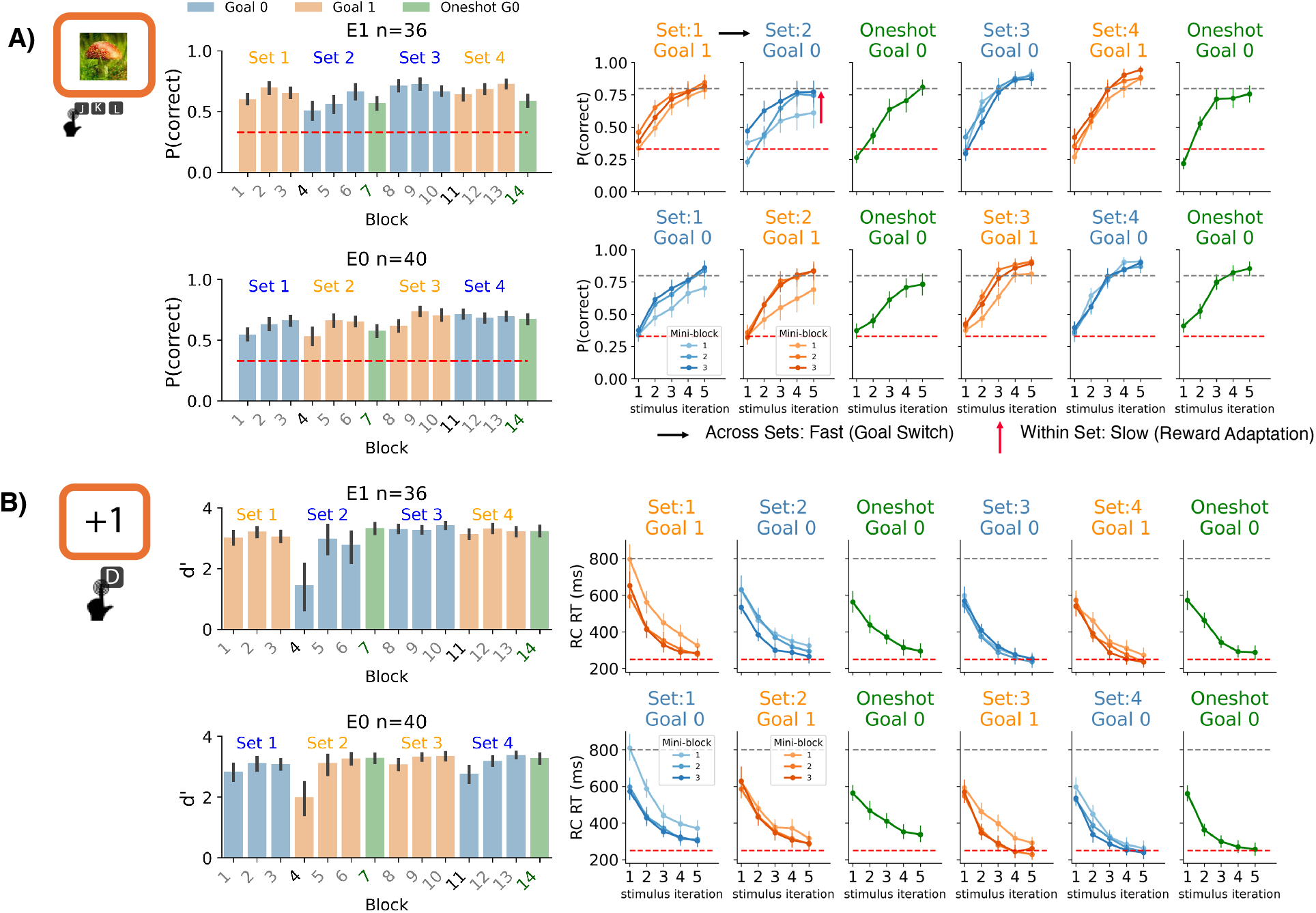
Experiment 2 behavioral performance. **A)** Stimulus-action learning performance. Left: Block-level mean choice accuracy across groups. Right: Mean accuracy across stimulus iterations. Error bars: 95% CI. Red dashed line: chance level (33.3%); Grey dashed line: 80% accuracy. **B)** Reward collection (RC) performance. Left: Block-level RC accuracy (d^′^) across groups. Right: Median RC reaction time across stimulus iterations. Error bars: 95% CI. Red dashed line: 250 ms; Grey dashed line: 800 ms.

Together, these findings indicate that slow valuation continues to enhance performance in the later stages of learning. By establishing stable, goal-congruent outcome–reward associations, extended practice complements fast goal switching, ultimately mitigating goal-switch costs across both learning and one-shot blocks.

#### Outcome processing accuracy improves with practice

Having shown that practice mitigates initial learning inefficiencies under Goal 0 and goal-switch cost in stimulus–action mappings, we next asked whether this pattern reflects a systematic change in outcome processing efficiency, measured by reward collection (RC). We first examined the goal-switch sets (Set 2 and 4) to determine whether switching to Goal 0 incurs a higher cost (Fig. 3B left). An LMM predicting each block’s RC *d*^′^ showed significant main effects of block set (*b* = 0.57, *SE* = 0.10, *t* = 5.52, *p* < .001, 95% CI [0.37, 0.78]; Supplementary Table 12), mini-block (*b* = 0.42, *SE* = 0.06, *t* = 6.54, *p* < .001, 95% CI [0.29, 0.54]), and two-way interaction between block set and mini-blocks (*b* = −0.48, *SE* = 0.13, *t*= −3.74, *p* < .001, 95% CI [−0.72, −0.23]), revealing practice improved outcome processing in both sequence groups. Furthermore, a significant two-way interaction between block set and sequence group emerged (*b* = −0.50, *SE* = 0.21, *t* = −2.41, *p* = .016, 95% CI [−0.91, −0.09]). Follow-up simple-effects analyses showed that average RC accuracy across mini-blocks differed only in the early stage between groups, with significantly higher performance in the E0 group learning under Goal 1 over the E1 group learning under Goal 0 in Set 2 (Holm-adjusted pairwise comparisons of estimated marginal means: Δ = 0.389, *SE* = 0.178, *t*(162) = 2.19, *p* = .030), but not in Set 4. Consistent with stimulus-action learning, the results showed that outcome processing accuracy under Goal 0 also improved with practice.

Next, we tested within-group switch costs in outcome processing accuracy (*d*^′^). Indeed, early in learning (Set 1 to Set 2), switching goals led to significantly lower *d*^′^ in both the E1 group (Goal 1 to Goal 0: *M* = 3.54 vs. 2.65; two-sided paired *t*-test: *t*(35) = 2.96, *p* = .006, 95% CI [0.28, 1.50], *d* = 0.66, BF_10_ = 7.06) and the E0 group (Goal 0 to Goal 1: *M* = 3.49 vs. 3.15; *t*(39) = 2.07, *p* = .045, 95% CI [0.01, 0.68], *d* = 0.37, BF_10_ = 1.17). However, unlike stimulus-action learning, where late-stage switch costs were absent, switching from Set 3 to Set 4 still lowered *d*^′^ for the E1 group (Goal 0 to Goal 1: *M* = 4.01 vs. 3.77; *t*(35) = 2.51, *p* = .017, 95% CI [0.05, 0.43], *d* = 0.56, BF_10_ = 2.71), but not for the E0 group (Goal 1 to Goal 0: *M* = 3.77 vs. 3.59; *t*(39) = 1.78, *p* = .083, 95% CI [−0.03, 0.39], *d* = 0.32, BF_10_ = 0.71), with no significant difference in the switch cost between the groups. This residual cost in the E1 group may reflect the preceding one-shot Goal 0 block (Block 7), which reinforced the Goal 0 representation and increased the cost of switching back to Goal 1.

Conversely, an LMM predicting outcome processing speed (RC RT) from block set, mini-block, and sequence group revealed significant main effects only for block set (*b* = −0.32, *SE* = 0.04, *t* = −8.92, *p* < .001, 95% CI [−0.39, −0.25]) and mini-block (*b* = −0.16, *SE* = 0.02, *t* = −7.11, *p* < .001, 95% CI [−0.20, −0.11]; Supplementary Table 13), with no significant main effect of sequence group or interactions (*p*s ≥ .074), reflecting a general practice-induced acceleration in response execution across both groups. Finally, we evaluated within-group changes in the RC RT difference across goal switches. Across both early (Set 1 to Set 2) and late (Set 3 to Set 4) goal switches, we found no significant change (*p*s ≥ .586, BF_10_ ≤ 0.20) across switches, except that the E0 group exhibited a significant acceleration when switching from Goal 0 to Goal 1 in the early stage (*M* = 0.01 vs. −0.13; *t*(39) = 2.26, *p* = .030, 95% CI [0.01, 0.26], *d* = 0.45, BF_10_ = 1.64, two-sided paired *t*-test).

Together, these findings demonstrate that extended practice improves both the accuracy and speed of goal-congruent outcome evaluation, especially in overcoming the initial cost of processing Goal 0 outcomes; this improvement raises the critical question of whether outcome processing directly drives the efficiency of stimulus–action learning.

#### Alignment of internal and instructed goals predicts learning efficiency

To test whether accurate outcome valuation supports stimulus–action learning, we next examined how participants’ trial-by-trial outcome processing related to their choice performance. We hypothesized that an individual’s outcome collection behavior reflects their internal goals (e.g., either collecting outcome +1 or withholding collection for outcome +0 was classified as having Goal 1 as the internal goal). To formalize this, we quantified goal alignment as “instructed goal occupancy” — defined as the proportion of trials on which a participant’s inferred internal goal matched the instructed goal of the block. A three-way GLMM predicting choice accuracy revealed no significant interaction among instructed goal occupancy, goal-switch block (Block 4 vs. 11), and stimulus iteration. However, instructed goal occupancy interacted significantly with stimulus iteration (*β* = 3.888, *SE* = 0.0.541, *z* = 7.183, *p* < .001; Supplementary Table 14), indicating that higher goal occupancy was associated with steeper learning curves. This aligns with the expectation that synchronizing internal representations with instructed goals facilitates learning valid stimulus–action–outcome associations.

#### Goal alignment predicts faster outcome processing with practice

We next asked whether this goal alignment also predicts the speed of outcome processing. An LMM predicting block-level RC RT from instructed goal occupancy (inferred from outcome key presses), goal switch sets (Set 2 vs. 4), and stimulus-action learning accuracy, showed that RC RT was significantly faster with higher instructed/correct goal proportion (*β* = −1.036, *SE* = 0.265, *t* = −3.911, *p* < .001), higher accuracy (*β* = −1.397, *SE* = 0.161, *t* = −8.658, *p* < .001) and in later block sets (*β* = −0.186, *SE* = 0.041, *t* = −4.528, *p* < .001; Supplementary Table 15).

Although there was no three-way interaction, instructed goal occupancy interacted reliably with block set (*β* = −1.493, *SE* = 0.527, *t* = −2.833, *p* = .005). Interestingly, follow-up estimated marginal means showed that RC RT decreased more strongly across block sets only among participants with high instructed-goal occupancy (*b* = −0.467, *SE* = 0.087, *t*(442.35) = −5.360, *p* < .001), whereas no reliable change was observed among those with low instructed-goal occupancy. Taken together, these results show that with practice, greater goal alignment was associated with increasingly faster outcome processing, marking a transition from effortful goal reconfiguration early in learning toward more efficient goal-switching after extended practice.

### Cognitive model of goal-dependent valuation dynamics

The behavioral results in Experiment 2 suggest that goal-dependent learning involves both rapid alignment with instructed goals and slower adjustment of outcome-reward properties. We next used a reinforcement learning (RL) framework to determine whether these dynamics could be dissociated computationally and to characterize their contribution to individual differences in learning (Fig. 4A). Specifically, we aimed to examine whether the subjective outcome valuation is governed by two components: (1) a *fast component* reflecting executive control that switches goal representations across block sets, and (2) a *slow component* capturing the gradual updating of associations between outcomes and their reward properties (positive vs. negative, with initial association set by prior experience; operating within a block set). More modeling details can be found in the **Methods** section.

**Figure 4.**
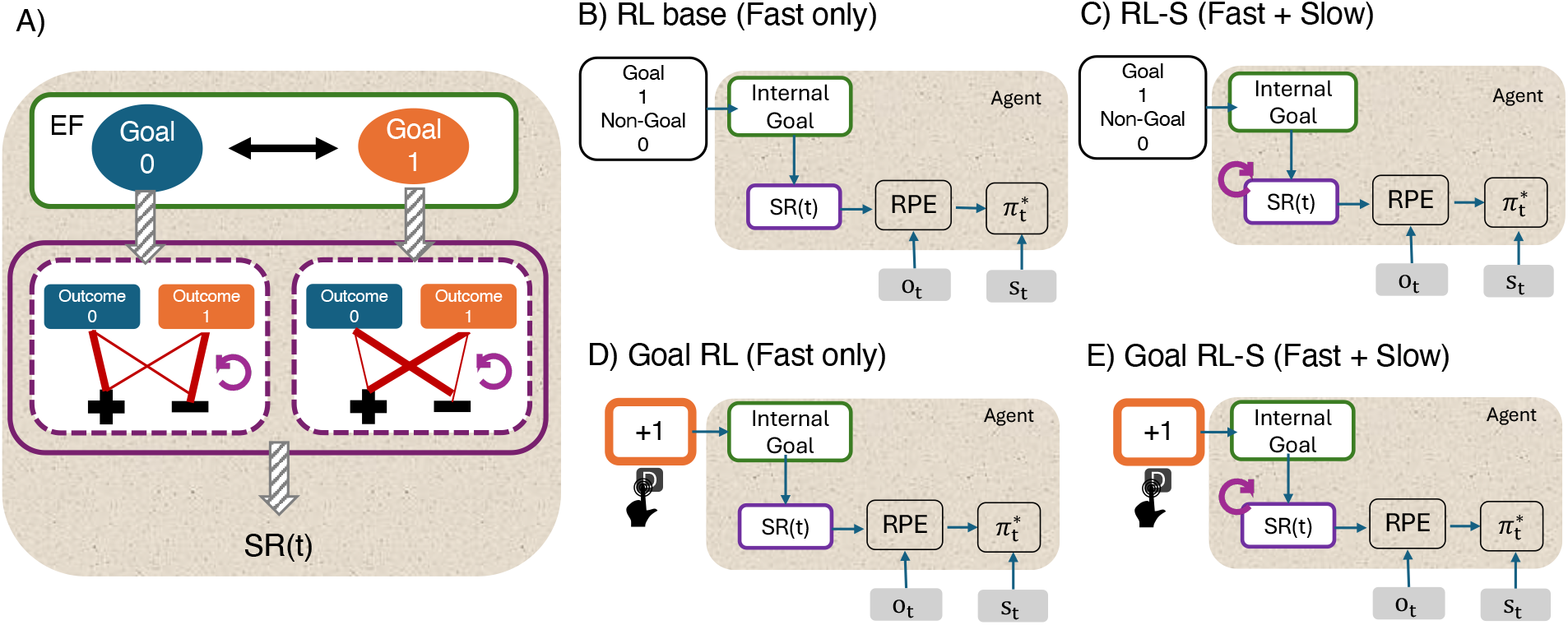
Computational modeling framework. **A)** Schema of the goal-dependent reinforcement learning (RL) model incorporating a fast goal switch and slow outcome-reward adaptation. SR(t) denotes the subjective reward on trial t. **B-D)** Four RL model variants applied to Experiment 2. On each trial t, the agent computes a reward prediction error (RPE) from the observed outcome o_t_ and updates its policy π^∗^ for the presented stimulus s_t_. **B)** RL base: Standard Q-learning model assuming a fast goal switch conditioned on instructed goals. **C)** RL-S: Model incorporating both a fast goal switch and a slow outcome-reward update conditioned on instructed goals. **D)** Goal RL: Model with a fast goal switch conditioned on internal goals inferred from reward collection key presses. **E)** Goal RL-S: Model incorporating both a fast goal switch and a slow outcome-reward update conditioned on internal goals. Note that goals were always instructed in this experiment. RL models imply perfect adherence to the instructions, while Goal RL models allow for imperfect alignment due to conflicting information from personal preferences or experience.

We developed four variants of Q-learning models that learn stimulus-action-outcome associations via trial and error: RL base, RL-S, Goal RL, and Goal RL-S (Fig. 4 B-E). These models differ along two primary dimensions:

- **Goal alignment**: RL base and RL-S assume perfect adherence to instructed goals. By contrast, Goal RL and Goal RL-S account for potential misalignment between instructed and internal goals, where we inferred internal goals from reward-collection key presses (i.e., an outcome is considered a goal if the participant attempted to collect it on that trial). Note that for a participant with perfect goal alignment, the Goal RL and base RL models were indistinguishable, as were the Goal RL-S and RL-S models.
- **Outcome-reward value adaptation**: RL base and Goal RL assume a one-shot update in outcome value, whereas RL-S and Goal RL-S incorporate a slow update in outcome-reward association. Note that the fast only models are nested in the -S models, with a slow update learning rate of 1.

At the group level, Goal RL-S provided the best overall account of the data, as indicated by the lowest Akaike Information Criterion (AIC; Fig. 5A). Simulations using fitted parameters revealed that models incorporating internal goals (Goal RL and Goal RL-S) were uniquely capable of capturing performance during the early switch block (Block 4; Fig. 5B), in which some participants had not yet attuned to the newly instructed goals. Goal RL-S additionally captured the gradual change in subjective outcome valuation observed across Blocks 4–6 (Fig. 5B).

**Figure 5.**
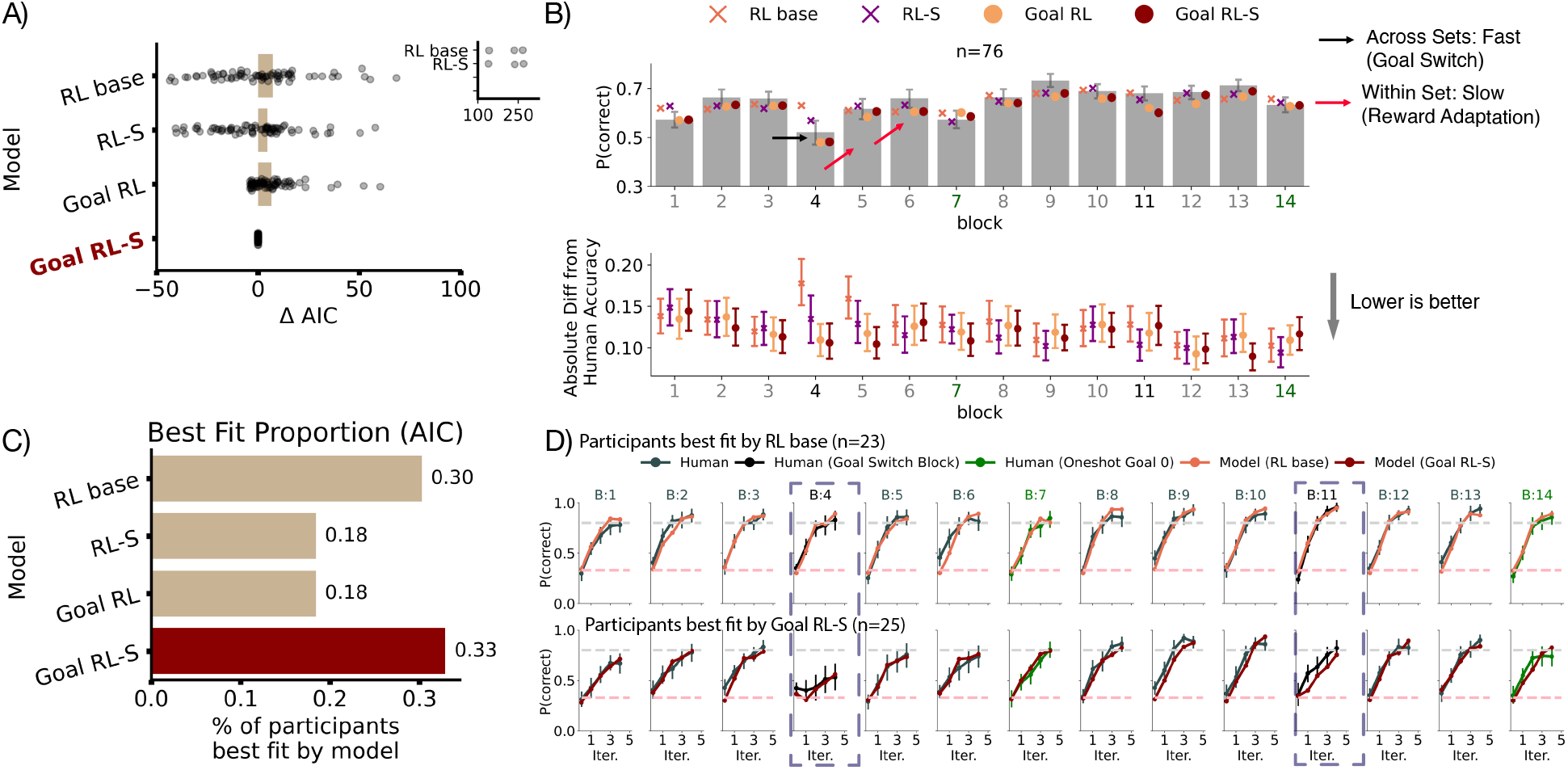
Model comparison and validation. **A)** Group-level model comparison based on average AIC. Values represent the ΔAIC relative to the winning model (Goal RL-S). The inset shows outlier participants for RL base and RL-S. **B)** Model validation against empirical data. Top: Block-level mean accuracy for human participants (bars) and model simulations (dots). Bottom: Absolute difference in block-level accuracy between each model and human data (lower is a better fit). **C)** Individual-level model comparison showing the proportion of participants best fit by each candidate model (lowest AIC). **D)** Empirical vs. simulated learning curves split by best-fitting model group. Top: RL base model simulations vs. participants’ best fit by RL base. Bottom: Goal RL-S model simulations vs. participants’ best fit by Goal RL-S. All model simulations in **B** and **D** were generated using individual best-fitting parameters. Error bars: 95% CI.

**Figure 6.**
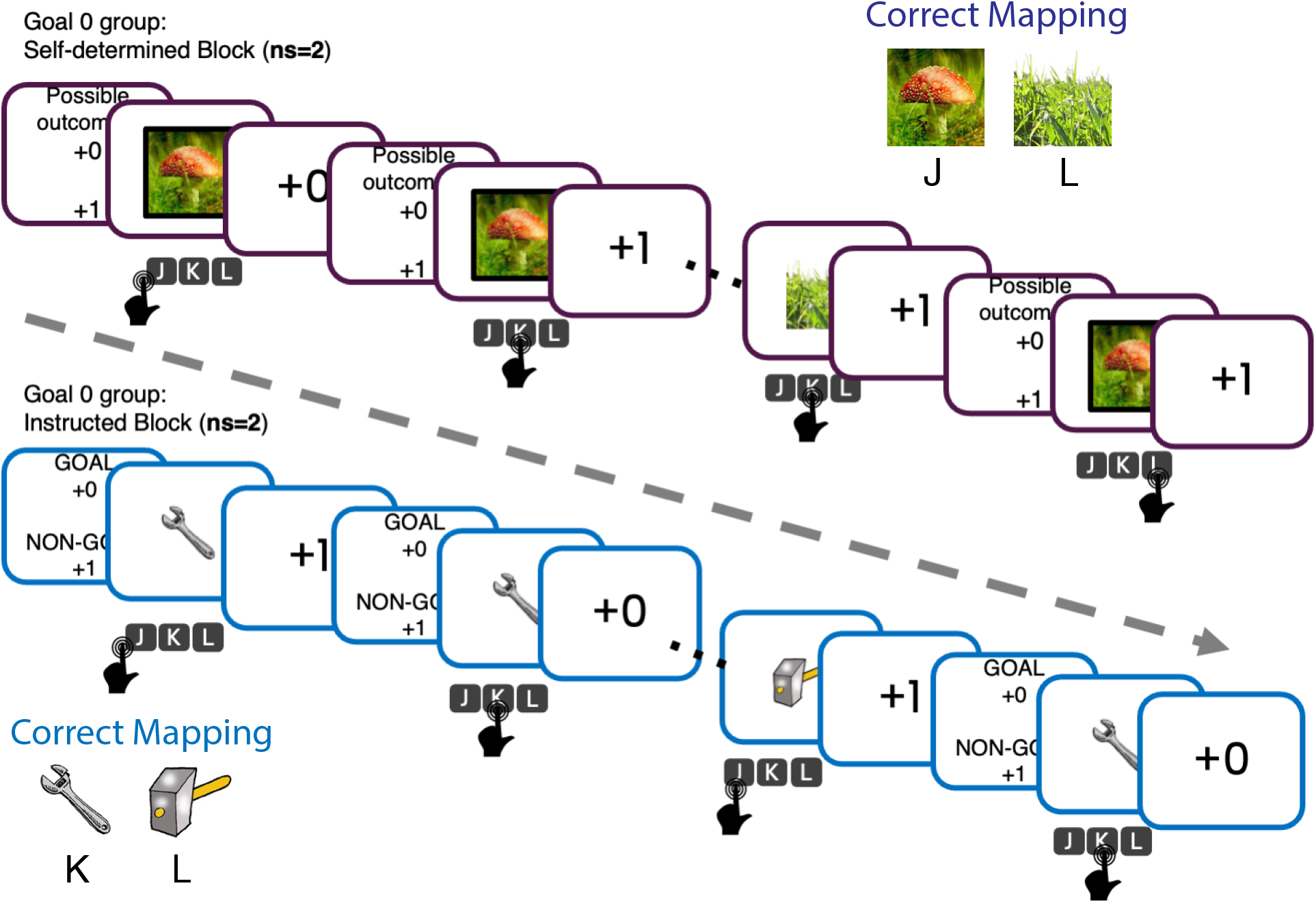
Example: Goal 0 group in Experiment 1 Prior to the first block (the self-determined block), participants viewed the following instructions: “The game’s rules have been lost. Please try to figure out how to play on your own by pressing keys after each image and observing the outcomes. [Press SPACEBAR to continue]”. During this self-determined block, participants in the Goal 0 condition frequently obtained Outcome 1 because two keys yielded this outcome for each stimulus. Upon transitioning to the second block (the instructed block), participants were told: “Good News. The rules of the games

**Figure 7.**
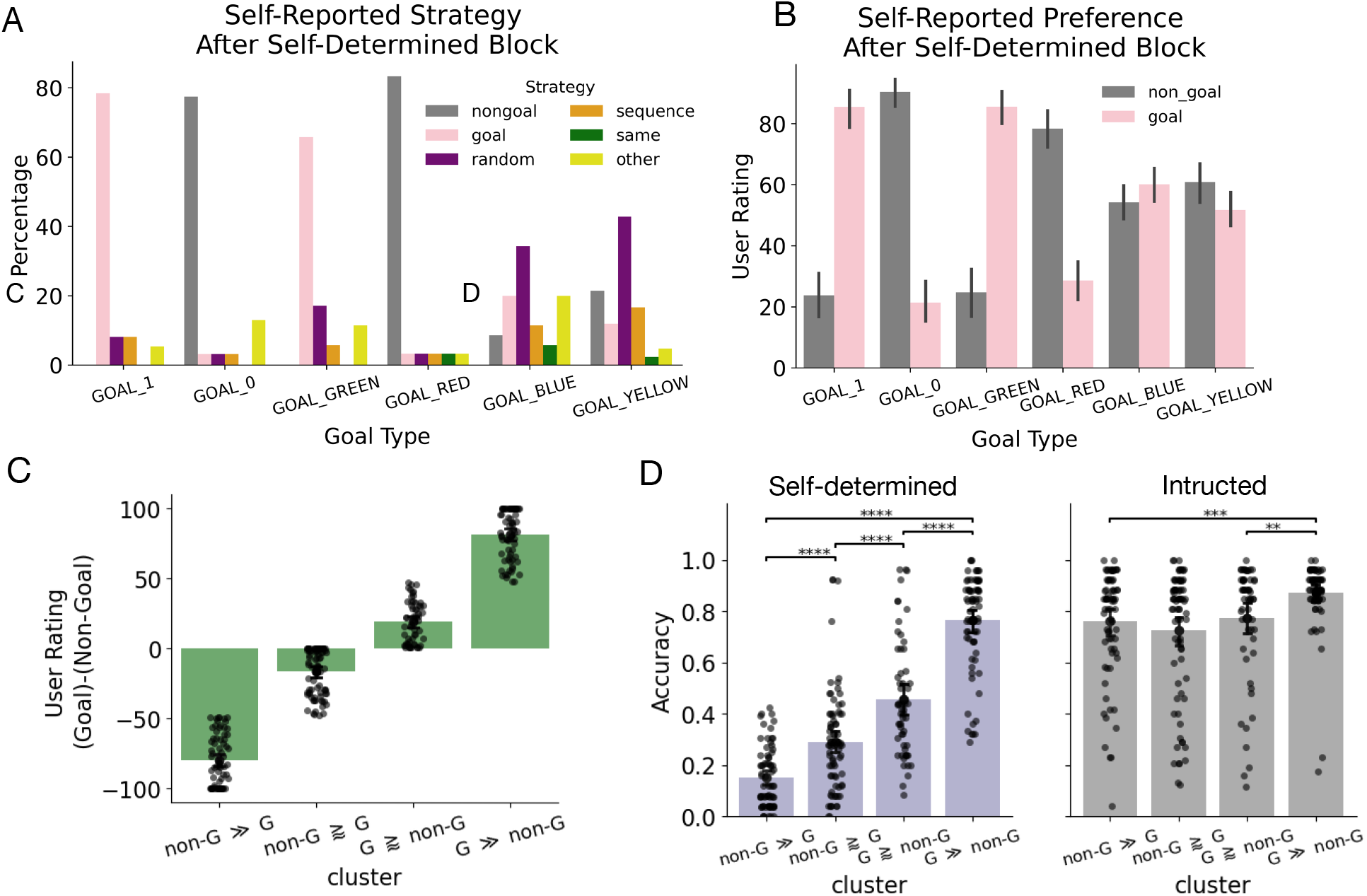
Self-report questionnaire in Exp. 1 **A)** Self-reported strategies. The sample questionnaire in the Goal 0 group is as follows. Goal: Collect as many outcome 0 as possible; Non-goal: Collect as many outcome 1 as possible; Random: Choose keys randomly without any specific pattern. *Sequence*: Use a specific sequence of keys regardless of outcome. Same: Always press the same key throughout the game. Others: An open text box for reporting alternative strategies. **B)** Self-reported preference. Participants rated how much they like each outcome on a scale from 1 to 10. **C)** Quantile-based cluster (equal-sized buckets) of outcome preference (user rating). **D)** Preference cluster against accuracy in different blocks. Error bar: 95% CI. ^∗∗^*p <* 0.01; ^∗∗∗^*p <* 0.001; ^∗∗∗∗^*p <* 0.0001*; Unpaired two-sided t-test with Benjamini–Hochberg correction for multiple comparisons*.

However, individual-level model comparison revealed participant heterogeneity: RL base and Goal RL-S each best accounted for roughly one-third of the sample (Fig. 5C). Specifically, participants best fit by RL base maintained consistently high performance across all blocks with one-shot reward-outcome property update. By contrast, participants best fit by Goal RL-S struggled to adapt immediately during goal-switch blocks (Blocks 4 and 11) and exhibited a slow, continuous outcome-reward updating process throughout the switch block sets (Blocks 4–6 and 11–13; Fig. 5D). Across the switch block sets, the two groups differed significantly in accuracy only in the first switch block (pairwise two-sided *t*-tests, Block 4: *t*(42.85) = 1.61, *p*_adj_ = .006, *d* = 0.46). Although accuracy was also lower for Goal RL-S in Block 11 (*t*(42.18) = 2.21, *p* = .033, *d* = 0.63), this difference did not survive correction (*p*_adj_ = .172, BF10 = 1.99). No group differences emerged in subsequent blocks (all *p*adj ≥ .099), suggesting that participants varied in their initial susceptibility to prior outcome-reward associations, but this difference diminished with extended practice. All model validation can be found in Supplementary Fig. 9& 10.

#### The influence of slow adaptation on learning

To validate whether the model-derived slow updating parameter meaningfully relates to learning performance, we first sought to confirm that individual estimates of *γ* (update rate of subjective reward) predicted learning accuracy.

Among participants best characterized by models incorporating slow outcome-reward updating (RL-S and Goal RL-S; *n* = 39), an LMM predicting choice accuracy showed no three-way interaction between goal-switch block set (Block set 2 vs. 4), mini blocks, and model parameter *γ* of the instructed goal. Nonetheless, a significant positive main effect of *γ* on mean accuracy indicated that faster updates were associated with higher accuracy (*β* = 0.445*, SE*= 0.097*, t*= 4.580, *p* < .001; Supplementary Table 16). In addition, a significant negative interaction between *γ* and mini-block (*β* = −0.123*, SE*= 0.044*, t*= −2.813*, p* = .005) indicated that the effect of *γ* on accuracy was more pronounced in the first mini-block (early switch) than the last mini-block (late switch) in goal-switch block sets. This pattern suggests that differences in outcome-reward updating were most consequential early in learning, when the new goal–outcome mappings were being established.

Having confirmed that *γ* predicted learning accuracy, we next asked whether individual differences in *γ* also related to outcome processing speed, given prior evidence linking faster outcome processing to better learning performance (Chen et al., 2026; Molinaro & Collins, 2026). An LMM predicting outcome processing speed, as indexed by RC RT, showed a main effect of *γ* negatively predicting RC RT (*β* = −0.556, *SE* = 0.262, *t* = −2.127, *p* = .033; Supplementary Table 17). This finding suggested that participants with higher *γ* values exhibited faster outcome processing. In addition, the main effects of block set (*β* = −0.343, *SE* = 0.131, *t* = −2.627, *p* = .009) and mini-block (*β* = −0.161, *SE* = 0.030, *t* = −5.344, *p* < .001) indicated that outcome processing sped up as the task progressed (practice effect).

In summary, rapid alignment with the instructed goal supported flexible adaptation following goal switches, but initially incurred a switch cost that diminished with practice. In parallel, the rate of outcome-reward updating (*γ*) modulated learning across both early and late stages and was positively associated with accelerated outcome processing. Together, these results suggest that a fast component governs the flexible updating of goal representations, whereas a slow component stabilizes value representations to support automated execution.

## Discussion

Human goal-dependent learning exhibits great flexibility. Beyond adapting to abstract and novel goals (McDougle et al., 2022; Molinaro & Collins, 2023), our results showed that this flexibility extends to counterintuitive goals (pursuing +0 over +1), consistent with prior work (Pan et al., 2025; Shi et al., 2026). However, this flexibility in fast goal-dependent valuation is not perfect, and increases with more exposure to the same goal. Indeed, we found that initial learning was less efficient under Goal 0 than under Goal 1, suggesting that overriding the default tendency to pursue standard rewarding outcomes (+1) imposes additional cognitive demands. In addition to this fast flexible process allowing us to rapidly learn from changing goals, we additionally observed slower dynamics. Specifically, repeatedly pursuing the same goal improved performance; in particular, it attenuated the initial learning inefficiency under the counterintuitive Goal 0.

Consistent with the existence of a slow process that strengthens our valuation of a repeatedly pursued goal, we found that goal switching incurred a substantial cost on learning, regardless of switching to Goal 0 or Goal 1. The cost of switching to Goal 1 was particularly striking given participants’ strong default tendency to pursue Goal 1 in Experiment 1: three consecutive blocks of pursuing Goal 0 led to a strong impairment in subsequently learning from Goal 1. While switching costs are well documented in the task-switching literature, where changing tasks or response rules typically impairs performance (Monsell, 2003; Robbins, 2007), our findings differ from typical rule switching. Specifically, in our case, the goal switch doesn’t impact the rules the participant follows to make stimulus-action choices (as those are learned); instead, it impacts how outcomes are interpreted. We showed that switching the goal that guides outcome evaluation impairs goal-directed learning.

In addition to a fast goal valuation process, enabling rapid new goal pursuit, and a slower one, enabling adjustment across repetitions of the same block, we additionally found evidence for slower dynamics across the whole task, whereby effects in the first half of the experiment decreased or disappeared in the second part. In particular, the effect of Goal 0 vs. Goal 1 largely (though not completely) disappeared. Similarly, the learning cost incurred after a goal switch was markedly reduced and absent at later stages of learning. Goal 0 evaluation and Goal switching therefore became more efficient with experience, even when the newly instructed goal conflicted with the previously established one. This practice-driven improvement reflects a slow establishment of stable, goal-congruent outcome–reward associations that complement the rapid goal-switch process; crucially, in a form of meta-learning, participants became more effective at retrieving previously learned outcome-reward associations from Goal cues.

This practice-dependent stabilization of goal representations was also corroborated by outcome processing speed (measured via RC RT). Participants with high goal alignment (a strong match between their internal and instructed goals)) exhibited a pronounced acceleration in RC RT over time, whereas those with low goal alignment showed no reliable change. In line with prior work in goal-dependent automaticity (Verplanken & Aarts, 1999; Aarts & Dijksterhuis, 2000), this suggests that practice strengthens efficient binding between current goals and their corresponding outcome-reward mappings, enabling even initially slow-updating individuals to flexibly adapt.

To dissect the cognitive mechanisms underlying this phenomenon, we developed a computational framework that incorporates complementary fast and slow valuation processes to account for goal-dependent learning under changing and opposing goals. Although a model integrating both processes best captured group-level behavior, our model comparison revealed substantial heterogeneity in individual goal-dependent valuation. This heterogeneity reflects varying efficiencies in the fast and slow processes, leading to differences in individuals’ susceptibility to prior experience. For instance, some learners exhibited rapid goal-switching that quickly aligned with instructed goals (best captured by the RL-based and RL-S models), while some were effective at forming goal-congruent outcome-reward associations (best captured by the RL-based and Goal RL models). Crucially, an initial susceptibility to prior experience did not impose a permanent limit on performance, as early inefficiencies can be compensated for by the slower learning mechanism. Consequently, two individuals may ultimately achieve equally successful goal-dependent learning, even if they differ substantially in how much prior experience influenced their behavior during early practice.

Furthermore, our finding of stabilization in goal-dependent learning aligns closely with prefrontal-striatal modulation frameworks from neuroscience studies (Terra et al., 2020; Gore et al., 2023). Koszeghy and Passecker (2026) posited a dual mechanism underlying goal-directed behavior: fast switching via direct matrix projections and slower, learning-based adaptation via a striosome-dopamine pathway. In early learning, goal switching necessitates slower adaptation via prefrontal cortex (PFC)-modulated striosomal compartments to establish new goal-dependent outcome-reward associations. Following extended practice, the PFC can exploit direct matrix projections to execute rapid switching over these stabilized mappings. This physiological distinction accounts for late-stage learning dynamics, where most participants switch effectively between opposing goals regardless of initial learning discrepancies. Future neuroimaging and electrophysiological studies could leverage this experimental paradigm and computational modeling framework to delineate the precise mechanisms by which the PFC coordinates both striosome-mediated learning and direct matrix projections.

Our computational framework also provides a principled approach to disambiguate whether initial goal adaptation failures stem from poor goal-switch control (Goal RL), outcome sensitivity (RL-S), or a combination of both (Goal RL-S). The behavioral heterogeneity captured by our model converges with evidence that working memory constraints (limiting goal-switch control) (Molinaro & Collins, 2023) or outcome sensitivity (intrinsic outcome preference) can affect outcome valuation (Zhu et al., 2025; Shi et al., 2026). Future work combining this paradigm with secondary measures (e.g., working memory batteries or outcome-preference questionnaires) and clinical populations could test whether these computational phenotypes characterize meaningful individual differences in goal-dependent learning.

Finally, our observed practice-dependent attenuation of experience-driven biases parallels research on overcoming Pavlovian biases, which are traditionally viewed as automatic and hardwired. Although evidence for overwriting Pavlovian biases through cognitive training is mixed (Ereira et al., 2021; Schurr et al., 2024), Fleming et al. (2025) showed that practice reduces Pavlovian bias only when paired with high-conflict training. Our task design incorporated alternating, opposing goals, mirroring key elements of high-conflict regimes (albeit without a separate control condition). Crucially, our claim is limited to an attenuation of prior influences rather than the lasting bias reduction achieved through multi-day training. Because our experiment was restricted to a single one-hour session, it remains unclear whether these improvements would persist on subsequent days. Future research should systematically vary learning curricula and utilize multi-day practices to assess the durability of this effect.

In conclusion, our findings showed how complementary fast and slow goal-dependent valuation processes enable extended practice to overcome constraints imposed by prior experience or limited cognitive control, ultimately establishing flexible goal pursuit. Furthermore, the proposed novel computational framework allows future research to pinpoint distinct computational phenotypes across individuals by examining the dynamic interplay among multiple valuation mechanisms.

## Materials and Methods

### Behavioral Experiments

#### Participants

A total of 300 participants completed Experiment 1 online through Prolific (151 female; age: *M* = 27.8, *SD* = 6.11). We recruited 50 participants for each goal pair. Seventeen participants were excluded based on predefined criteria: excessive missing trials (more than 5 consecutive missing trials or 10 missing trials in a block), more than 15 repeated responses, and fast average response times (mean lower than 300ms), yielding a final sample of 283 participants. In both experiments, we applied the elbow method to identify the exclusion threshold.

In Experiment 2, a total of 126 participants participated online (73 female; age: *M* = 26.7, *SD* = 7.0): 54 were recruited from the undergraduate participant pool at the University of California, Berkeley, and 72 were recruited via Prolific. We excluded 47 participants who did not reach 50% accuracy across the practice and learning phases, and an additional three participants who failed to recognize the goal outcome on the questionnaire following one-shot blocks. The final sample consisted of 76 participants (*n* = 40 in sequence group E0; *n* = 36 in sequence group E1). In both experiments, all participants provided informed consent online. Undergraduates received partial course credit as compensation, while Prolific participants were compensated at a rate of $12 per hour (10 min in Experiment 1; 1 h in Experiment 2).

#### Procedure

All experiments started with an informed consent form and ended with a demographic survey. The study procedures were approved by the Institutional Review Board of the University of Berkeley. In Experiment 1, participants went through a brief introduction to the task, explaining that they needed to learn the associations between 2 stimuli and 3 actions by trial and error. Participants were then told that *“The game’s rules have been lost. Please try to figure out how to play on your own by pressing keys after each image and observing the outcomes.”* (Supplementary Fig. 6), followed by a self-determined block. Afterwards, they rated in a brief survey their preference for each outcome and the strategy that best describes the rule they used to match the images with keys in the game. Participants were then told that “The rules of the game have been found. Your job is to try to obtain a specific goal.” and then finished with an instructed block with an explicit instructed goal and non-goal. Experiment 1 comprised two blocks of 26 trials each (13 iterations × 2 stimuli). Each trial began with the possible outcome (1,800 ms), followed by stimulus presentation (1,500 ms), feedback (1,500 ms), and an inter-trial fixation cross (500 ms). On average, participants completed the task in approximately 10 minutes.

Experiment 2 began with a brief instruction of the task, explaining that they needed to learn the associations between 6 stimuli and 3 actions by trial and error and collect the goal outcome with a separate key press. Participants then went through a brief practice round, where they learned the associations between 2 stimuli and 3 actions with 13 iterations per stimulus. Both the instruction and practice round used a yellow and a blue solid circle as goal and non-goal outcomes, respectively, to avoid any prior practice on specific goal outcomes from influencing the main learning blocks. After the practice round, participants started the learning phase, where they experienced one of two goal sequences (Fig. 2 B). Participants were assigned to one of two sequence structures consisting of four 3-block sets interspersed with one-shot Goal 0 blocks at Blocks 7 and 14. The E1 (Early 1 Late 0) group completed sets in a Goal 1, 0, 0, and 1 order (Blocks 1–3, 4–6, 8–10, and 11–13, respectively). Conversely, the E0 group followed an inverted set sequence (Goals 0, 1, 1, and 0), with the one-shot Goal 0 blocks at Blocks 7 and 14. Crucially, in each block, participants needed to re-learn different pairs of stimulus-action mappings, ensuring that outcome processing can be dissociated from stimulus-action learning throughout the whole experiment. After each one-shot block, participants reported the correct goal outcome in the previous one-shot block in a short questionnaire. We excluded participants who answered incorrectly from further analysis (3 out of 126). Overall, Experiment 2 consisted of 14 learning blocks (5 iterations * 6 stimuli = 30 trials per block). Each trial began with the possible goal outcome (1,800 ms), followed by stimulus presentation (1,500 ms), feedback (1,500 ms), and an inter-trial fixation cross (500 ms). The experiment took around 1 hour to complete on average.

### Computational Modeling

All models are based on a reinforcement learning (RL) framework (Sutton & Barto, 2018). Specifically, we used a Q-learning model, which updates stimulus-action associations *Q* after observing stimuli, actions, and rewards (*s_t_, a_t_, r_t_*) at trial *t* through a classic delta rule:

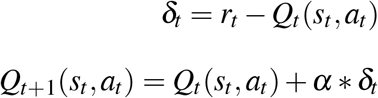

*Q*_0_ is initialized as <u>^1^</u>, where *n_A_* denotes the total number of possible actions in a task (three in Experiment 2). We used two learning rates *α* and *α*_oneshot_, denoting the learning rates in the learning and one-shot phases, respectively. Both are free parameters ∈ [0, 1]. The RL model employs a softmax policy with a fixed *β* = 20 to improve parameter recovery reliability (Wilson & Collins, 2019) (we verified in this context that including a free beta did not improve fit, as has been observed in other experiments with deterministic feedback (Pan et al., 2025; Senta et al., 2025)):

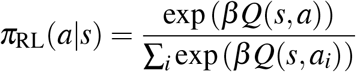

A“sticky” choice parameter *κ* ∈ [−1, 1] is also used to account for stimulus-independent choice perseveration – the tendency to repeat the same key press across consecutive trials regardless of the presented stimulus. This was integrated into the softmax policy as:

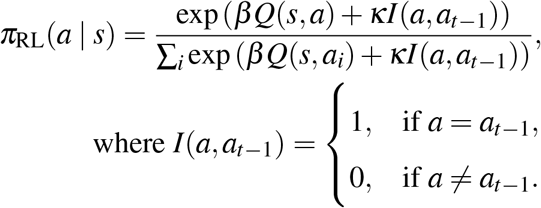

Finally, a random lapse in choices is added to the policy with a noise parameter *ε* ∈ [0, 1]:

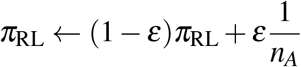

We have two *ε*: *ε*_early_ for blocks 1 to 7 and *ε*_late_ for blocks 8 to 14, to account for an observed practice effect that reduces choice stochasticity.

Below, we detail the four computational models evaluated in this study, beginning with the baseline model.

#### RL base

The base model follows the delta learning rule and choice policy described above. On each trial *t*, the reward prediction error *δ_t_*is computed from a subjective reward *SR_t_*, which is determined by whether the observed outcome *o_t_* matches the instructed goal *g*^instructed^ of the current block *b*:

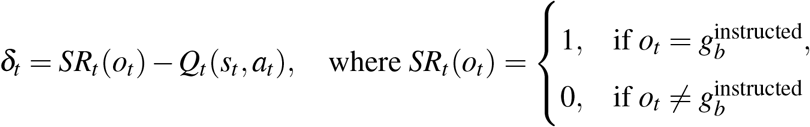

RL base thus consists of five free parameters: *α*, *α*_oneshot_,*κ*, *ε*_early_ and *ε*_late_.

#### RL-S

Building on top of the RL base model, RL-S incorporates a slow update of the subjective reward SR used to compute the reward prediction error. This update itself follows a delta rule:

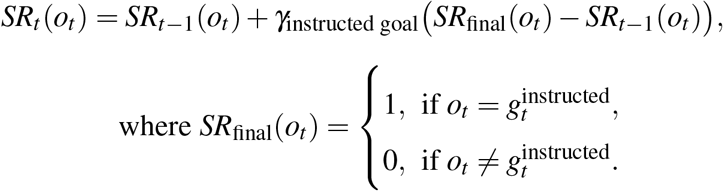

where *γ* _instructed goal_ is an update rate indexed by instructed goal, assuming that there exist different update rates for Goal 0 and Goal 1. *SR*_final_ represents the target value of the update. RL-S thus consists of seven free parameters: *α*, *α*_oneshot_, *κ*, *ε*_early_, *ε*_late_, *γ*_0_, and *γ*_1_

#### Goal RL

Similar to the base model, Goal RL follows the delta learning rule and choice policy. Here, we assume that the experienced subjective reward depends on internal goals rather than instructed goals. Suppose that *O* = {*o*_1_*, o*_2_} represents the set of possible outcomes (e.g., {0, 1}), where *o_t_* ∈ *O* is the outcome presented on trial *t*, and *o-_t_* represents the unpresented alternative. We inferred the internal goal *g*^internal^ via the outcome collection response *c_t_* ∈ {0, 1}.

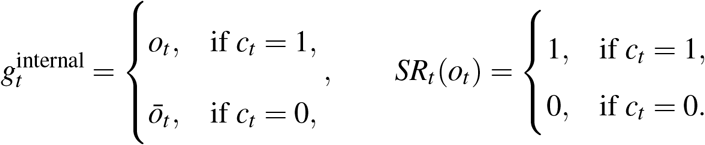

Therefore, collecting an outcome (*c_t_* = 1) indicates that the presented outcome matched the agent’s internal goal (*o_t_* = *g*^internal^), yielding *SR_t_* = 1. Conversely, withholding response (*c_t_* = 0) signifies a goal mismatch (*o_t_* = *g*^internal^), resulting in *SR_t_* = 0. Goal RL consists of five free parameters: *α*, *α*_oneshot_, *κ*, *ε*_early_, and *ε*_late_.

#### Goal RL-S

Similar to the RL-S model, Goal RL-S incorporates a slow reward property update, except that the update direction depends on internal goals:

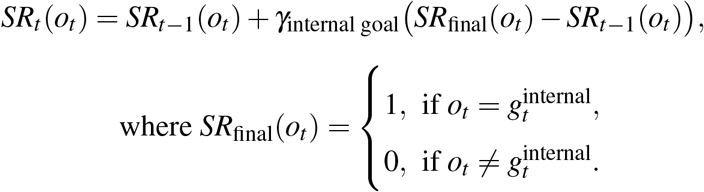

where the internal goal is derived in the same way as in the Goal RL model. Goal RL-S consists of seven free parameters: *α*, *α*_oneshot_, *κ*, *ε*_early_, *ε*_late_, *γ*_0_, and *γ*_1_

#### Fitting procedure

All models were fitted using maximum likelihood estimation with Python’s scipy library. The L-BFGS-B algorithm was applied for bound-constrained minimization using eight random starting points and a maximum of 1,000 iterations per participant (Byrd et al., 1995). Model identification was based on AIC metrics (Akaike, 1974). The distribution of fitting parameters and recovery results are in Supplementary Figs. 11&12.

### Statistical analysis

All group comparisons were conducted with an independent-samples Welch’s t-test. To quantify the magnitude of the difference, we calculated Hedges’ *g*, providing a standardized effect size that is robust to discrepancies in sample size. To assess whether participant accuracy significantly exceeded chance level, we performed one-sample, two-tailed t-tests for each goal type (within-group comparison). In all analyses, we controlled the false discovery rate (FDR) using the Benjamini-Hochberg procedure with an overall *α* level of 0.05. Finally, to quantify the strength of the evidence under a Bayesian framework, we computed the Bayes Factor (*BF*_10_). All statistical analyses were conducted in Python utilizing the pingouin and statsmodels libraries.

### Mixed-effects models

We used Generalized Linear Mixed-Effects Models (GLMMs) or Linear Mixed-Effects Models (LMMs) to assess the significance of fixed effects while controlling for participant-level variation with random intercepts. Mixed-effects models were fit using functions *lmer* and *glmer* from the R library *lme4*. All models were estimated using restricted maximum likelihood estimation (REML), while the likelihoods in GLMMs were estimated using Laplace approximation. In the case of significant interactions, we performed simple-effects analysis with pairwise comparisons of estimated marginal means and slopes (*emmeans*, *emtrends*). The mathematical formulations for models having simple-effects analyses are presented below, with all other model specifications provided in Supplementary .

#### Block-level RC performance trajectories across groups

An LMM predicts reward collection (RC) accuracy *d*^′^ as a function of the group, block set, mini block, and their interactions:

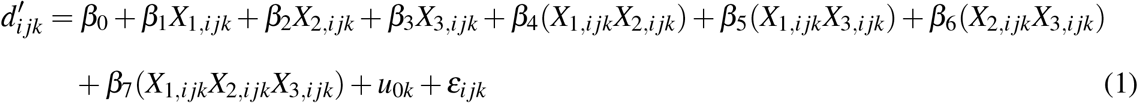

where:

- *d^’^_ijk_*: RC accuracy (*d*^′^) at mini-block *i* (1–3), block set *j* (2 vs. 4), for participant *k*.
- *X*_1*,i*_ *_jk_, X*_2*,i*_ *_jk_, X*_3*,i*_ *_jk_*: Predictors for group (*E*0 vs. *E*1), block set, and mini-block, respectively.
- *u*_0*k*_ ∼ *N* (0*, σ* ^2^): Participant-specific random intercept.
- *ε_i jk_* ∼ *N* (0*, σ* ^2^): Residual error.

In the follow-up simple analysis, the slopes representing mini-block learning rates across group were parameterized as:

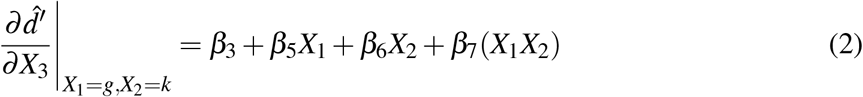

| Task Stage ($I_{\text{Set4}}$) | Group($X_1$) | Simple Slope Expression |
| --- | --- | --- |
| Block Set 2 ($I_{\text{Set4}} = 0$) | E0 ($g = 0$) | $\beta_3$ |
| | E1 ($g = 1$) | $\beta_3 + \beta_5$ |
| Block Set 4 ($I_{\text{Set4}} = 1$) | E0 ($g = 0$) | $\beta_3 + \beta_6$ |
| | E1 ($g = 1$) | $\beta_3 + \beta_5 + \beta_6 + \beta_7$ |

## Acknowledgments

This work was funded by award number 2336466 from the National Science Foundation.

T.P. is supported by the Berkeley Graduate Fellowship. We are grateful to Amitai Shenhav and Jason Leng for their feedback on the model design.

## Supplementary Information

### Experiment 1: Self-reported questionnaire and practice effect. Self-reported outcome preference partially explains the learning differences

In Experiment 1 (self-determined goal task), we observed stronger biases in numerical feedback compared to color feedback: when given explicit instructions, participants learned equally well for Goal Red and Goal Green conditions. This equivalence may reflect greater individual differences in red/green preferences, despite red’s typical association with negative outcomes. These findings prompted us to investigate whether performance variations could be driven by individual outcome preferences (e.g., some participants preferring red over green or blue over yellow).

We categorized participants based on their outcome ratings collected immediately after the minimum instruction block, creating four quantiles according to their relative preferences:

1. Non-G ≫ G (n=74) - Strongly favoring non-goal over goal outcomes
2. Non-G ≳ G (n=80) - Slightly favoring non-goal over goal outcomes
3. G ≳ Non-G (n=58) - Slightly favoring goal over non-goal outcomes
4. G ≫ Non-G (n=71) - Strongly favoring goal over non-goal outcomes.

Results showed that participants who strongly favored goal outcomes (G ≫ Non-G) performed better than all other groups (F_(3,279)_ = 146.2, *p* < .001, BF*_incl_ >* 100). Importantly, 88.7% of participants in the G ≫ Non-G group were from Goal +1 and Goal Green types, suggesting that +1 and green carry strong positive associations from prior experience.

However, individual preferences alone cannot fully explain the observed learning differences. We found no evidence of performance differences between the G ≳ Non-G group and groups that favored non-goal outcomes (slightly favor non-goal: *t*(279) = 1.293, *p* = .568, BF_10_ = 0.332; strongly favor non-goal: *t*(279) = 0.378, *p* = .982, BF_10_ = 0.119). Notably, the G ≳ Non-G group consisted primarily of participants in Goal Yellow and Goal Blue types—outcomes that typically lack strong positive or negative associations. This pattern may reflect the influence of our task instructions, where participants were told to “figure out the lost rule.” This framing may have led participants to infer underlying design principles, making their learning dependent on objective prior associations rather than purely subjective preferences. Specifically, +1 and green are objectively associated with positivity in real-world contexts, while yellow and blue lack such clear associations.

### Practice effects on learning performance

To isolate the effects of practice between blocks, we conducted a control experiment using a Goal 1 task with two instructed blocks, where participants received complete instructions throughout the entire task. We compared this fully instructed (I-I) design with the Goal 1 group in the original self-determined + instructed (SD-I) design.

An LMM showed a three-way interaction effect among block (Block 1 vs. Block 2), task type (I-I vs. SD-I), and iteration (log-transformed) (*β* = 0.79 ± 0.29, *z* = 2.8, *p* < .01; Table 1). This suggests that the rate of learning varied depending on both task type and block. Specifically, block/practice effect alone cannot account for the behavioral patterns observed in our main SD-I task (Fig. 8 B). Even when pursuing the same outcome across blocks, participants still required cognitive effort to re-engage controlled processing when transitioning from self-determined to explicit instructions in the second block, which is also suggested by the strong evidence of the differences in response time between blocks in SD-I (BF_10_ = 56.6, with median posterior *δ* = −0.525, 95% CI [−0.835, −0.221]) but not in fully instructed (BF_10_ = 0.265, with median posterior *δ* = 0.143, 95% CI [−0.146, 0.434]). This implies that the instructional manipulation influences the observed learning differences, even in Goal 1.

**Figure 8.**
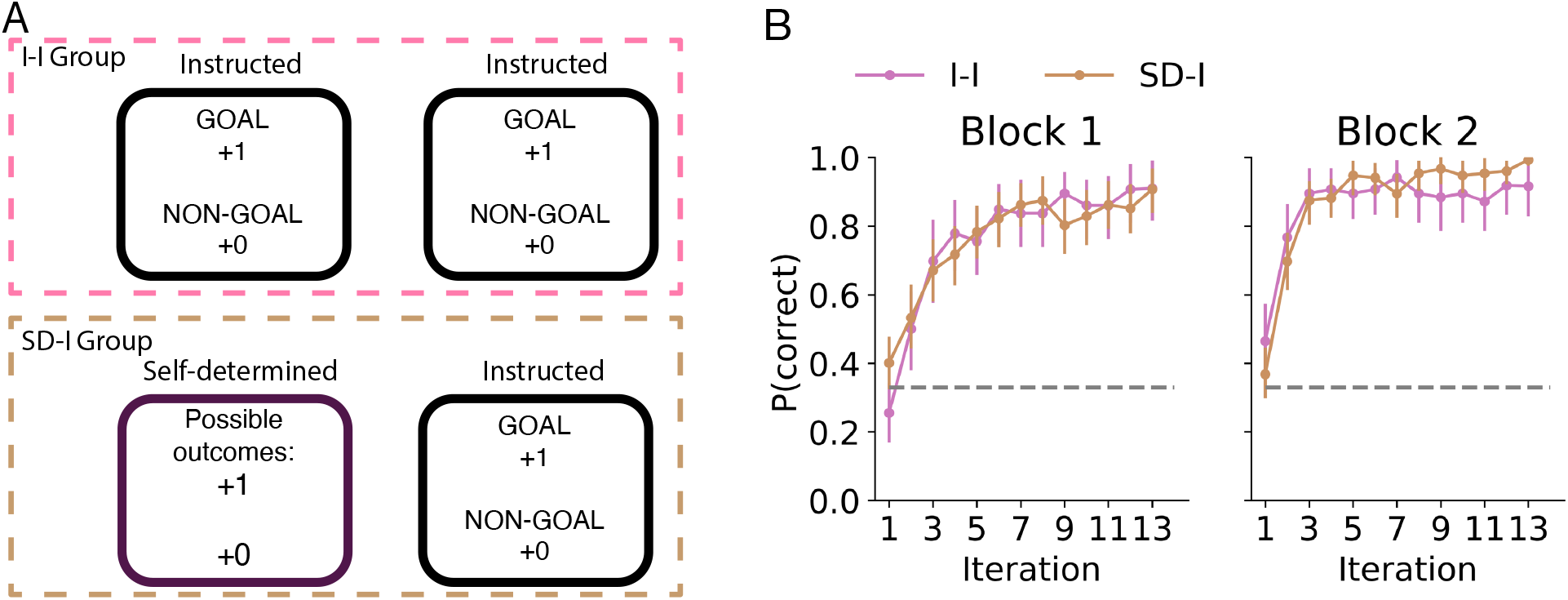
A) Task structure of fully Instructed (I-I) vs. Self-determined + Instructed task (SD-I). I-I: n=43. SD-I: n=46. **B)** Mean accuracy across stimulus iterations.

**Table 1.** GLMM: correct ∼ block * task type * log(stim_iter) + (block | id).

| Fixed effects: | Estimate | Std. Error | z value | p |
| --- | --- | --- | --- | --- |
| Intercept | -1.315 | 0.324 | -4.064 | < .001 |
| Block | 1.722 | 0.398 | 4.330 | < .001 |
| Task type | 0.487 | 0.435 | 1.121 | .262 |
| log(stim_iter) | 2.153 | 0.162 | 13.273 | < .001 |
| Block × Task type | -1.323 | 0.534 | -2.476 | .013 |
| Block × log(stim_iter) | -0.574 | 0.226 | -2.545 | .011 |
| Task type × log(stim_iter) | -0.633 | 0.201 | -3.153 | .002 |
| Block × Task type × log(stim_iter) | 0.793 | 0.287 | 2.764 | .006 |
| Random effects: | Variance | Std. Dev. | Corr. |  |
| Participant (id) |  |  |  |  |
| Intercept | 2.149 | 1.466 |  |  |
| Block | 2.475 | 1.573 | -0.763 |  |

**Figure 9.**
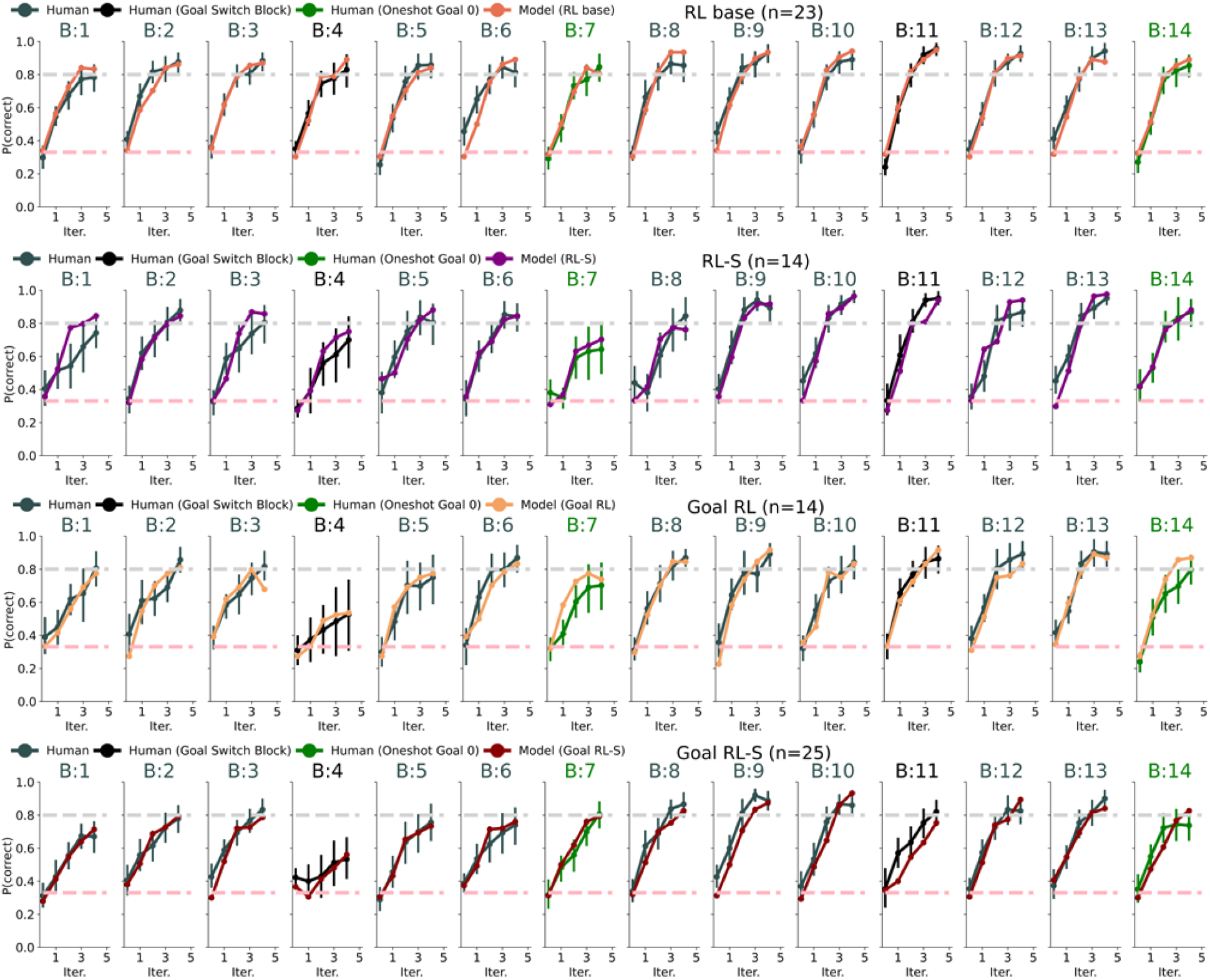
Best model validation for four models: Rows correspond to the subset of participants best captured by a given model based on AIC, and columns indicate different blocks; for each group (row), empirical behavior is compared to the corresponding best-fitting model.

**Figure 10.**
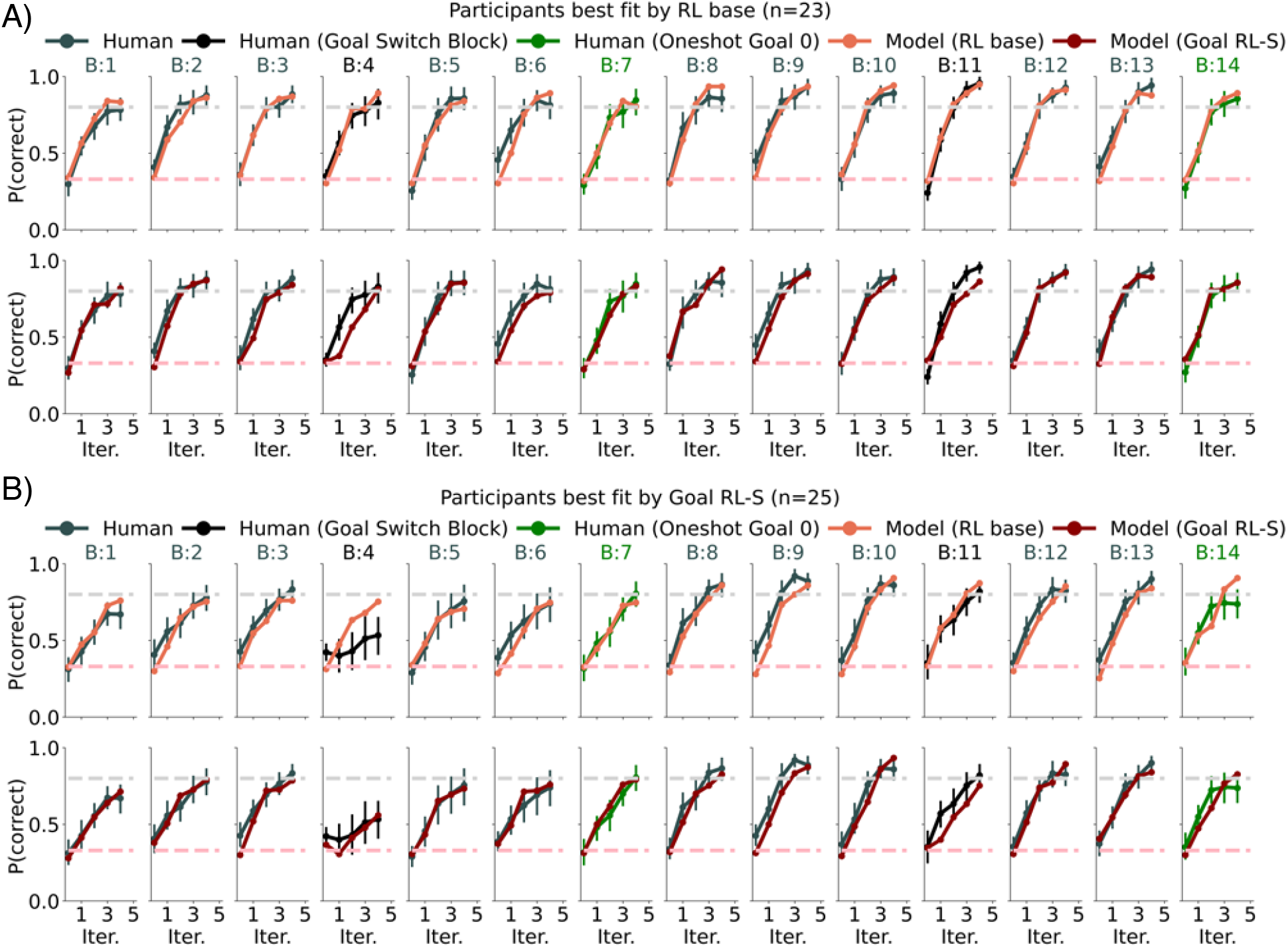
Contrast model validation: Rows correspond to the model simulation from RL base and Goal *RL-S models. Columns indicate different blocks. A) Participants best captured by RL base model. B) Participants best captured by Goal RL-S model. Notably, RL base model fails to capture the significant performance drop in Block 4 (the early goal-switch block)*.

**Figure 11.**
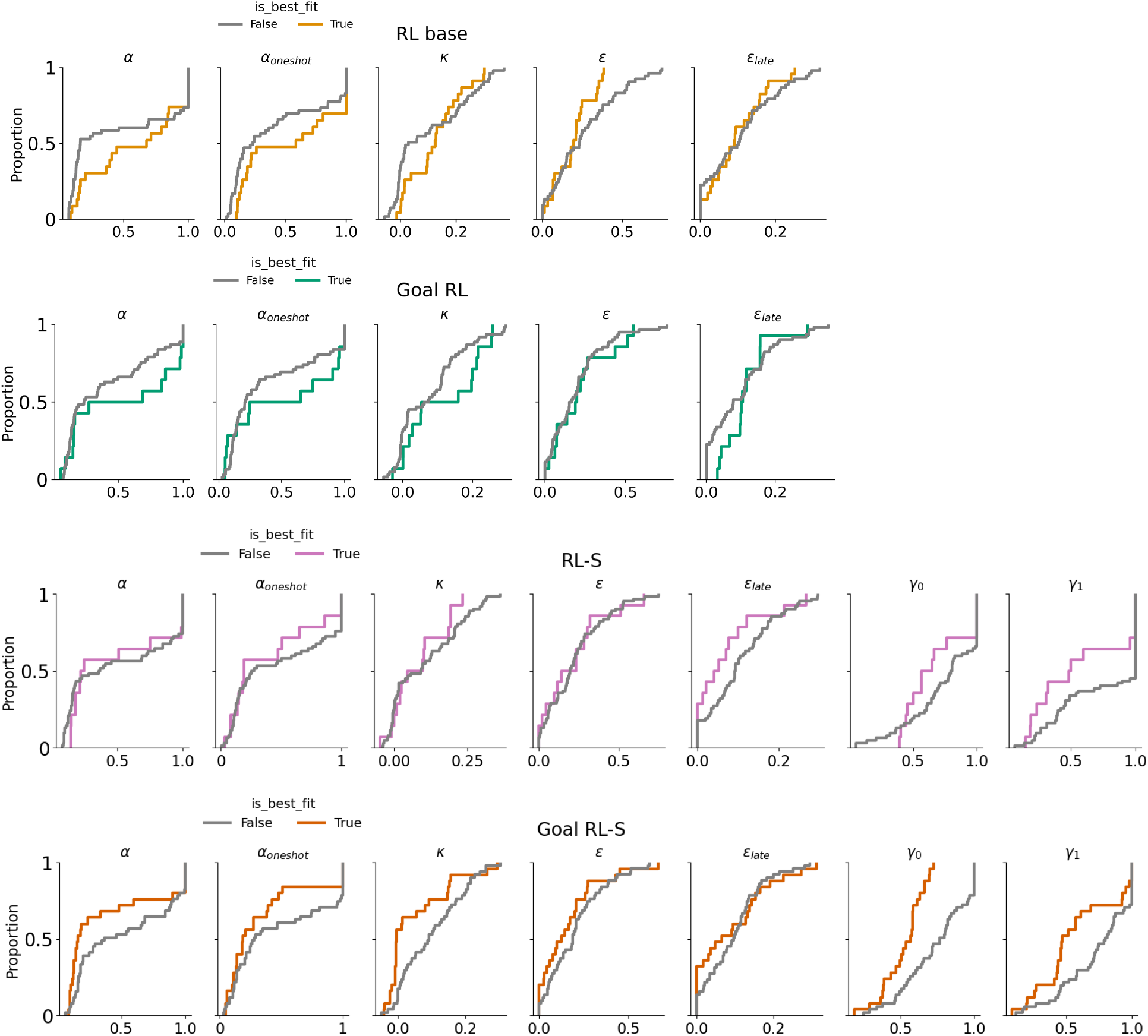
Fitted parameters across all cognitive models, experiment 2. Rows represent distinct models, and columns represent individual parameters. The x-axis: the value of the parameters. The y-axis: the proportion of participants with parameter value below y. Grey lines: participants not best fit by the model; colored lines: participants best fit by the model.

**Figure 12.**
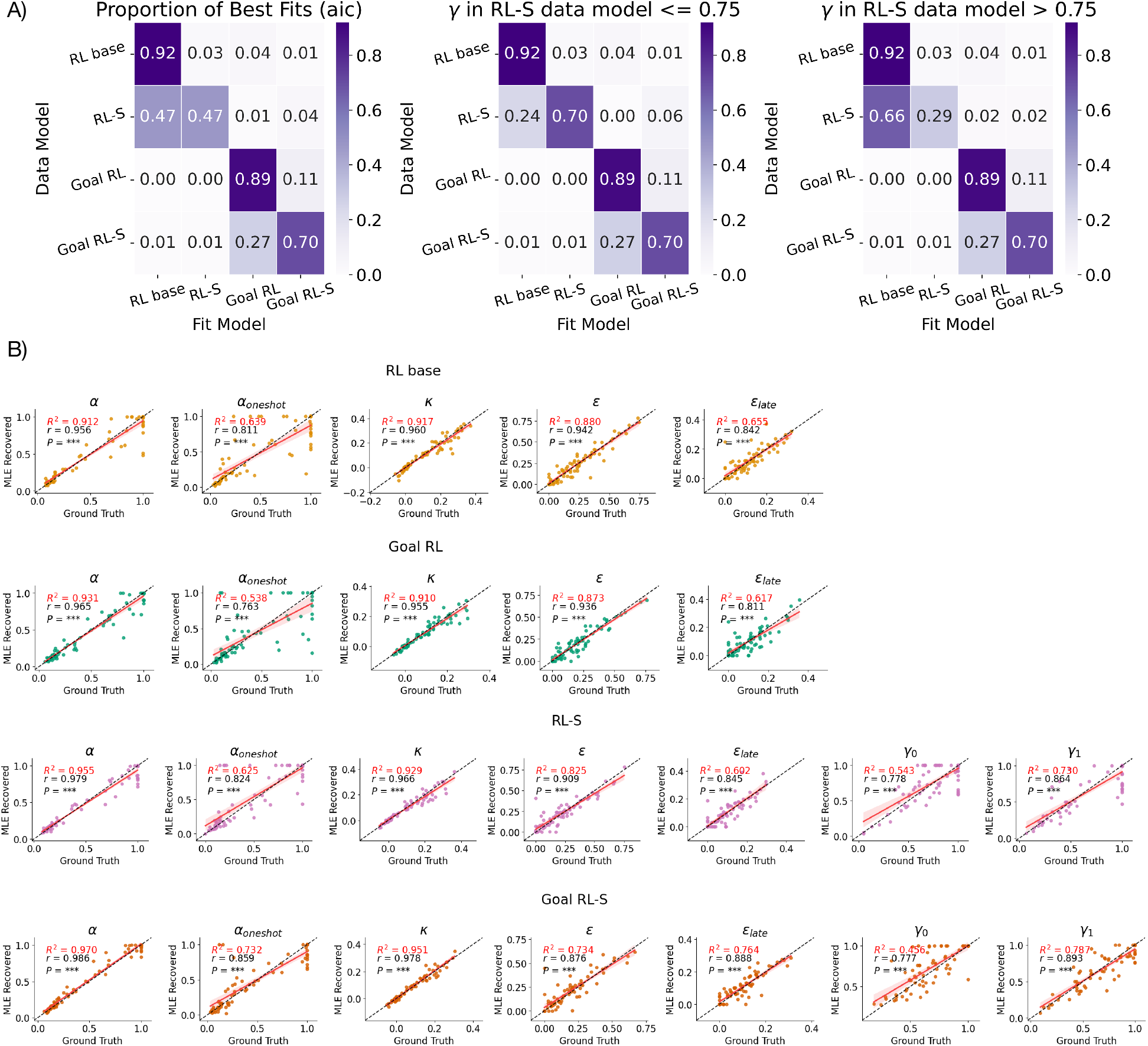
(A) Model identification All synthetic data were simulated from fitted human parameters. As anticipated, in the higher γ range, the RL-S model is less identifiable than the RL base. Right: confusion matrix for all models; Middle: confusion matrix for low γ in RL-S; Left: confusion matrix for high γ in RL-S. **(B) Parameter recovery** Recovered parameters exhibit high correlation with true parameters (p < 0.001). Rows represent distinct models, and columns represent individual parameters. Details of GLMM/LMM results

## Details of GLMM/LMM results

**Table 2.** LMM: correct ∼ block * counter * pair_type + (1 | id), where block is either self-determined or instructed, counter is true for Goal 0 and Goal Red, and false for Goal 1 and Goal Green. Pair type is color pair vs. numerical pair.

| Fixed effects: | Estimate | Std. Error | *t* value | *p* |
| --- | --- | --- | --- | --- |
| Intercept | 0.614 | 0.011 | 56.346 | < .001 |
| Block | −0.358 | 0.015 | −23.850 | < .001 |
| Counter | 0.306 | 0.022 | 14.051 | < .001 |
| Pair type | 0.014 | 0.022 | 0.663 | .508 |
| Block × Counter | 0.493 | 0.030 | 16.446 | < .001 |
| Block × Pair type | 0.003 | 0.030 | 0.115 | .909 |
| Counter × Pair type | 0.094 | 0.044 | 2.158 | .032 |
| Block × Counter × Pair type | 0.060 | 0.060 | 0.993 | .321 |
| Random effects: | Variance | Std. Dev. |  |  |
| Participant (id) | 0.012 | 0.109 |  |  |
| Residual | 0.042 | 0.206 |  |  |

*Note.* Statistically significant *p*-values ( $p < .05$ ) are shown in bold.

**Table 3.** GLMM results for Block 1: correct ∼ seq_group * log_stim_iter_c + (1 | id), where log_stim_iter_c is the centered, log-transformed stimulus iteration. seq_group was effect-coded (E1 = −0.5, E0 = +0.5).

| Fixed effects: | Estimate | Std. Error | z value | p |
| --- | --- | --- | --- | --- |
| Intercept | 1.029 | 0.077 | 13.438 | < .001 |
| Sequence Group | -1.200 | 0.104 | -11.510 | < .001 |
| Stimulus Iteration | 1.059 | 0.061 | 17.391 | < .001 |
| Sequence Group × Stimulus Iteration | -0.157 | 0.121 | -1.291 | .197 |
| Random effects: | Variance | Std. Dev. |  |  |
| Participant (id) | 0.279 | 0.528 |  |  |

**Table 4.** GLMM results for Block 4: correct ∼ seq_group * log_stim_iter + (1 | id), where *log_stim_iter is the log-transformed stimulus iteration. seq_group coded E1/E0 as −0.5/+0.5*.

| Fixed effects: | Estimate | Std. Error | z value | p |
| --- | --- | --- | --- | --- |
| Intercept | 0.029 | 0.125 | 0.229 | .819 |
| Sequence Group | 0.092 | 0.251 | 0.367 | .713 |
| Stimulus Iteration | 0.918 | 0.086 | 10.722 | < .001 |
| Sequence Group × Stimulus Iteration | 0.365 | 0.170 | 2.144 | .032 |
| Random effects: | Variance | Std. Dev. |  |  |
| Participant (id) | 1.016 | 1.008 |  |  |

**Table 5.** LMM results for Set 1: correct ∼ seq_group * mini_block + (1 | id), where *mini_block is centered. seq_group coded E1/E0 as −0.5/+0.5*.

| Fixed effects: | Estimate | Std. Error | t value | p |
| --- | --- | --- | --- | --- |
| Intercept | 0.632 | 0.011 | 57.287 | < .001 |
| Sequence Group | -0.037 | 0.022 | -1.681 | .093 |
| Mini-block | 0.042 | 0.010 | 4.307 | < .001 |
| Sequence Group × Mini-block | 0.032 | 0.020 | 1.628 | .104 |
| Random effects: | Variance | Std. Dev. |  |  |
| Participant (id) | 0.004 | 0.066 |  |  |
| Residual | 0.015 | 0.121 |  |  |

**Table 6.**
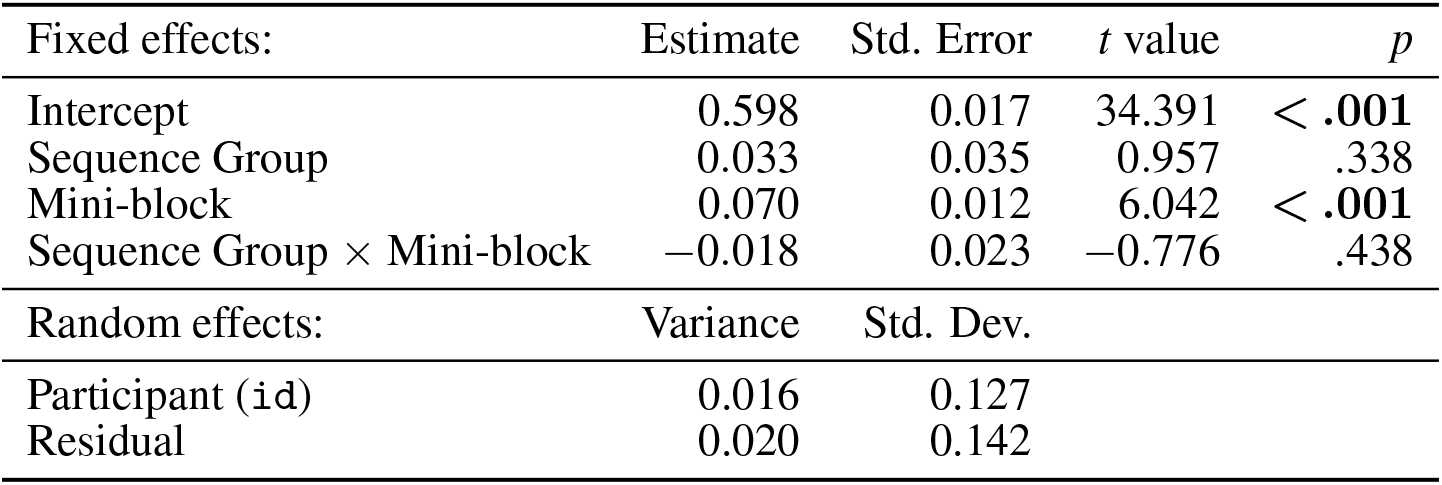
LMM results for Set 2: correct ∼ seq_group * mini_block + (1 | id), where *mini_block is centered. seq_group coded E1/E0 as −0.5/+0.5*.

| Fixed effects: | Estimate | Std. Error | *t* value | *p* |
| --- | --- | --- | --- | --- |
| Intercept | 0.598 | 0.017 | 34.391 | < .001 |
| Sequence Group | 0.033 | 0.035 | 0.957 | .338 |
| Mini-block | 0.070 | 0.012 | 6.042 | < .001 |
| Sequence Group × Mini-block | -0.018 | 0.023 | -0.776 | .438 |
| Random effects: | Variance | Std. Dev. |  |  |
| Participant (id) | 0.016 | 0.127 |  |  |
| Residual | 0.020 | 0.142 |  |  |

**Table 7.** GLMM results for block 8: correct ∼ seq_group * log_stim_iter + (1 | id), where *log_stim_iter is the log-transformed stimulus iteration. seq_group coded E1/E0 as −0.5/+0.5*.

| Fixed effects: | Estimate | Std. Error | *z* value | *p* |
| --- | --- | --- | --- | --- |
| Intercept | 0.854 | 0.091 | 9.385 | < .001 |
| Sequence Group | -0.638 | 0.181 | -3.521 | < .001 |
| Stimulus Iteration | 1.699 | 0.094 | 17.994 | < .001 |
| Sequence Group × Stimulus Iteration | -0.584 | 0.185 | -3.150 | .002 |
| Random effects: | Variance | Std. Dev. |  |  |
| Participant (id) | 0.408 | 0.639 |  |  |

**Table 8.** GLMM results for Block 11: correct ∼ seq_group * log_stim_iter + (1 | id), where *log_stim_iter is the log-transformed stimulus iteration. seq_group coded E1/E0 as −0.5/+0.5*.

| Fixed effects: | Estimate | Std. Error | *z* value | *p* |
| --- | --- | --- | --- | --- |
| Intercept | 0.942 | 0.091 | 10.405 | < .001 |
| Sequence Group | 0.457 | 0.180 | 2.540 | .011 |
| Stimulus Iteration | 1.946 | 0.098 | 19.845 | < .001 |
| Sequence Group × Stimulus Iteration | 0.059 | 0.192 | 0.309 | .758 |
| Random effects: | Variance | Std. Dev. |  |  |
| Participant (id) | 0.392 | 0.626 |  |  |

**Table 9.** LMM results for Set 4: accuracy ∼ seq_group * mini_block + (1 | id), where *mini_block is centered. seq_group coded E1/E0 as −0.5/+0.5*.

| Fixed effects: | Estimate | Std. Error | *t* value | *p* |
| --- | --- | --- | --- | --- |
| Intercept | 0.692 | 0.009 | 73.708 | < .001 |
| Sequence Group | 0.012 | 0.019 | 0.622 | .534 |
| Mini-block | 0.018 | 0.008 | 2.123 | .034 |
| Sequence Group × Mini-block | -0.051 | 0.017 | -3.033 | .002 |
| Random effects: | Variance | Std. Dev. |  |  |
| Participant (id) | 0.003 | 0.055 |  |  |
| Residual | 0.011 | 0.104 |  |  |

**Table 10.**
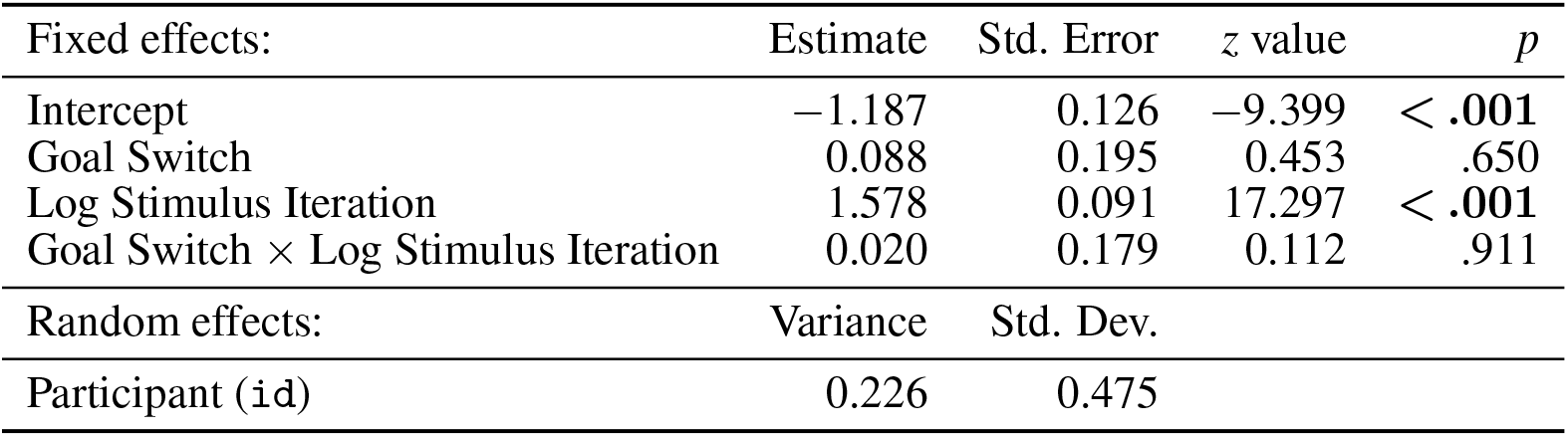
GLMM in one-shot blocks from E1 group: correct ∼ has_goal_switch * log_stim_iter *+ (1 | id), where has_goal_switch is coded no-switch as −0.5 and switch as 0.5, and log_stim_iter is the log-transformed stimulus iteration*.

**Table 11.** GLMM in one-shot blocks from E0 group: correct ∼ has_goal_switch * log_stim_iter *+ (1 | id), where has_goal_switch is coded no-switch as −0.5 and switch as 0.5, and log_stim_iter is the log-transformed stimulus repetition count*.

| Fixed effects: | Estimate | Std. Error | *z* value | *p* |
| --- | --- | --- | --- | --- |
| Intercept | -0.654 | 0.111 | -5.892 | < .001 |
| Goal Switch | -0.136 | 0.172 | -0.794 | .427 |
| Log Stimulus Iteration | 1.250 | 0.082 | 15.270 | < .001 |
| Goal Switch × Log Stimulus Iteration | -0.375 | 0.162 | -2.315 | .021 |
| Random effects: | Variance | Std. Dev. |  |  |
| Participant (id) | 0.196 | 0.442 |  |  |

**Table 12.** LMM results for Set 2 and Set 4: d_prime ∼ mini_block * block_set * seq_group + *(1 | id), where mini_block is centered. seq_group and block_set sum-coded as −0.5/+0.5, mini_block is mean centered, and seq_group coded E1/E0 as −0.5/+0.5*.

| Fixed effects: | Estimate | Std. Error | *t* value | *p* |
| --- | --- | --- | --- | --- |
| Intercept | 2.888 | 0.072 | 40.042 | < .001 |
| Block Set | 0.573 | 0.104 | 5.524 | < .001 |
| Sequence Group | 0.139 | 0.144 | 0.963 | .336 |
| Mini-block | 0.415 | 0.063 | 6.541 | < .001 |
| Block Set × Sequence Group | -0.500 | 0.207 | -2.410 | .016 |
| Block Set × Mini-block | -0.475 | 0.127 | -3.743 | < .001 |
| Sequence Group × Mini-block | 0.114 | 0.127 | 0.899 | .369 |
| Block Set × Sequence Group × Mini-block | 0.287 | 0.254 | 1.128 | .259 |
| Random effects: | Variance | Std. Dev. |  |  |
| Participant (id) | 0.190 | 0.436 |  |  |
| Residual | 1.222 | 1.105 |  |  |

**Table 13.** LMM results for Set 2 and Set 4: rt ∼ seq_group * mini_block_c * block_set + (1 | id), where mini_block_c is centered. Both seq_group (E1 = −0.5, E0 = +0.5) and block_set (Set 2 = −0.5, Set 4 = +0.5) were effect-coded.

| Fixed effects: | Estimate | Std. Error | *t* value | *p* |
| --- | --- | --- | --- | --- |
| Intercept | -0.267 | 0.020 | -13.291 | < .001 |
| Block Set | -0.319 | 0.036 | -8.922 | < .001 |
| Sequence Group | -0.006 | 0.040 | -0.161 | .872 |
| Mini-block | -0.156 | 0.022 | -7.111 | < .001 |
| Block Set × Sequence Group | -0.104 | 0.072 | -1.447 | .148 |
| Block Set × Mini-block | 0.078 | 0.044 | 1.788 | .074 |
| Sequence Group × Mini-block | 0.063 | 0.044 | 1.438 | .150 |
| Block Set × Sequence Group × Mini-block | -0.001 | 0.088 | -0.009 | .992 |
| Random effects: | Variance | Std. Dev. |  |  |
| Participant (id) | 0.006 | 0.080 |  |  |
| Residual | 0.146 | 0.382 |  |  |

**Table 14.** GLMM: correct ∼ log_stim_iter * goal_prop_centered * block + (1 | id), where goal_prop_centered is mean-centered instructed goal occupancy, log_stim_iter is the log-transformed stimulus iteration, and block codes blocks 4 and 11 as −0.5/0.5.

| Fixed effects: | Estimate | Std. Error | z value | p |
| --- | --- | --- | --- | --- |
| Intercept | -0.751 | 0.097 | -7.747 | < .001 |
| Goal Occupancy (centered) | -0.642 | 0.621 | -1.033 | .302 |
| Block | -0.066 | 0.168 | -0.392 | .695 |
| log(Stimulus Iteration) | 1.287 | 0.077 | 16.817 | < .001 |
| Goal Occupancy × Block | -0.838 | 1.234 | -0.679 | .497 |
| Goal Occupancy × log(Stimulus Iteration) | 3.888 | 0.541 | 7.183 | < .001 |
| Block × log(Stimulus Iteration) | 0.455 | 0.152 | 2.992 | .003 |
| Goal Occupancy × Block × log(Stimulus Iteration) | 1.592 | 1.079 | 1.476 | .140 |
| Random effects: | Variance | Std. Dev. |  |  |
| Participant (id) | 0.185 | 0.430 |  |  |

**Table 15.** LMM: reward_collection_rt_z ∼ correct_within * block_set * goal_prop_centered + (1 | id), where goal_prop_centered is mean-centered instructed goal proportion, correct_within is block-level choice accuracy normalized by *within-participant mean, and block_set codes block sets 2 and 4 as −0.5/0.5*.

| Fixed effects: | Estimate | Std. Error | t value | p |
| --- | --- | --- | --- | --- |
| Intercept | -0.198 | 0.021 | -9.320 | < .001 |
| Goal Proportion | -1.036 | 0.265 | -3.911 | < .001 |
| Choice Accuracy | -1.397 | 0.161 | -8.658 | < .001 |
| Block Set | -0.186 | 0.041 | -4.528 | < .001 |
| Goal Proportion × Choice Accuracy | -4.236 | 1.666 | -2.542 | .011 |
| Goal Proportion × Block Set | -1.493 | 0.527 | -2.833 | .005 |
| Choice Accuracy × Block Set | 0.868 | 0.324 | 2.680 | .007 |
| Goal Proportion × Choice Accuracy × Block Set | -0.878 | 3.329 | -0.264 | .792 |
| Random effects: | Variance | Std. Dev. |  |  |
| Participant (id) | 0.002 | 0.046 |  |  |
| Residual | 0.125 | 0.354 |  |  |

**Table 16.** LMM: correct ∼ mini_block * block_set * gamma_centered + (1 | id), where gamma_centered is the mean-centered discount factor (γ), mini_block is mean-centered, and block_set codes block sets 2 and 4 as −0.5/0.5.

| Fixed effects: | Estimate | Std. Error | *t* value | *p* |
| --- | --- | --- | --- | --- |
| Intercept | 0.531 | 0.027 | 19.527 | < .001 |
| Mini-block | 0.046 | 0.011 | 4.187 | < .001 |
| Block Set | 0.203 | 0.048 | 4.250 | < .001 |
| $\gamma$ (centered) | 0.445 | 0.097 | 4.580 | < .001 |
| Mini-block × Block Set | -0.049 | 0.022 | -2.218 | .027 |
| Mini-block × $\gamma$ (centered) | -0.123 | 0.044 | -2.813 | .005 |
| Block Set × $\gamma$ (centered) | -0.097 | 0.199 | -0.490 | .624 |
| Mini-block × Block Set × $\gamma$ (centered) | -0.010 | 0.088 | -0.110 | .912 |
| Random effects: | Variance | Std. Dev. |  |  |
| Participant (id) | 0.007 | 0.082 |  |  |
| Residual | 0.019 | 0.138 |  |  |

**Table 17.** LMM: reward_collection_rt_z ∼ mini_block * block_set * gamma_centered + (1 | id), where gamma_centered is the mean-centered γ derived from cognitive models, mini_block is mean-centered, and block_set codes block sets 2 and 4 as −0.5/0.5.

| Fixed effects: | Estimate | Std. Error | *t* value | *p* |
| --- | --- | --- | --- | --- |
| Intercept | 0.061 | 0.067 | 0.905 | .365 |
| Mini-block | -0.161 | 0.030 | -5.344 | < .001 |
| Block Set | -0.343 | 0.131 | -2.627 | .009 |
| $\gamma$ (centered) | -0.556 | 0.262 | -2.127 | .033 |
| Mini-block × Block Set | 0.019 | 0.060 | 0.308 | .758 |
| Mini-block × $\gamma$ (centered) | 0.137 | 0.120 | 1.141 | .254 |
| Block Set × $\gamma$ (centered) | -0.903 | 0.526 | -1.716 | .086 |
| Mini-block × Block Set × $\gamma$ (centered) | 0.405 | 0.240 | 1.686 | .092 |
| Random effects: | Variance | Std. Dev. |  |  |
| Participant (id) | 0.009 | 0.095 |  |  |
| Residual | 0.142 | 0.377 |  |  |

## Notes

### Competing Interest Statement

The authors have declared no competing interest.

